# A Charge Detection Mass Spectrometer for the Analysis of Megadalton-sized Molecules

**DOI:** 10.64898/2026.05.28.728353

**Authors:** Jakub Ujma, Chris Wheeldon, Alistair Schofield, Michael Danby, David Eatough, David Bruton, Anisha Haris, Keith Richardson, David Langridge, Andy Jarrell, Jeffery M. Brown, Benjamin E. Draper, Martin F. Jarrold, Kevin Giles

**Author notes:** **Corresponding Author**, Jakub Ujma - Waters Corporation, Stamford Avenue, Altrincham Road, Wilmslow SK9 4AX, U.K.

## Abstract

Advances in Electrostatic Linear Ion Trap (ELIT) Charge Detection Mass Spectrometry (CDMS) over the past 10 years have revolutionized its use for analyzing very high–molecular-weight species such as protein complexes, viral vectors, vaccines, viruses, and amyloid fibrils. Nonetheless, ELIT-based CDMS has remained confined to a small number of specialized instrumentation groups, predominantly in academia, where large and complex home-built instruments are operated by highly skilled scientists in dedicated facilities. In this report, we discuss the primary challenges addressed in the design of a benchtop ELIT-based CDMS instrument. We highlight key design aspects of the hardware, acquisition modes, and control software, and we present important performance metrics (mass range, resolution, and sensitivity) demonstrated using samples representative of the technology’s key application areas.

## Introduction

Analysis of large molecules by mass spectrometry (MS) began when Dole first coupled electrospray^1^ ionization (ESI) to mass spectrometry in the 1960s to measure polymers.^2^ Interest in this technique exploded as Fenn and Gall showed that biological molecules such as peptides and proteins could be ionized intact for measurement by MS.^3–5^ The use of MS-compatible volatile buffers (most commonly ammonium acetate) to maintain protein complexes in their native stoichiometry emerged soon after the initial demonstrations of ESI-MS, and has since become foundational to what is now known as the “native mass spectrometry” approach (native MS).^6–9^

In the ESI of large analytes, gas-phase ions are thought to be formed via the charged residue model (CRM).^2^ Since charging in this model is stochastic, a distribution of multiply-charged ions is typically observed in the mass-to-charge (*m/z*) domain for analytes having more than a single ionizable site. Multiple strategies have been developed to deconvolute charge state distributions (CSDs) to determine the mass, however they all rely on at least partial resolution of individual peaks within a charge state distribution.^4,10–12^ In native MS, this is best achieved when the molecules are well desalted and gently desolvated, typically requiring the use of a specialised form of ESI known as nano-electrospray ionisation (nESI).^13–15^ In the conventional native MS workflows, practitioners tend to fabricate their own nESI emitters (via glass pulling and sputter coating). Samples are then manually loaded, and the applied voltage and emitter position is subsequently adjusted for optimum results. Because emitters are single-use, this skill- and labour-intensive process must be repeated for every sample.^16^

Despite its low throughput, the native MS approach has seen rapid growth since its inception, driven by the analytical performance of MS instrumentation (e.g. *m/z* and mass range, sensitivity, resolution, dynamic range) as well as the versatility enabled by tandem MS and also ion mobility MS approaches.^17–21^ Unfortunately, as the size of molecules increases so too does their mass heterogeneity. This is due to both natural heterogeneity (post-translational modifications, sequence and subcomponent ratio variants) as well as adduction of non-volatile species.^22^ Charge multiplicity and mass heterogeneity ultimately lead to overlapping CSDs, posing challenges even for high-resolution MS and advanced deconvolution methods in the determination of mass values. This generally limits the application of native MS for molecules above ∼1 MDa, as well as smaller heterogenous assemblies and heavily-glycosylated proteins.^17,21,23,24^

The need for characterization of novel large modalities is accelerating. The biologics revolution began with the first monoclonal antibody-based drugs^25,26^ and has now progressed to larger, more complex molecules such as antibody-drug conjugates,^27^ heavily glycosylated proteins^24^ and biological polymers.^28^ Lipid nanoparticles and virus-like particles (LNPs, VLPs) are increasingly used in vaccines, while adeno-associated virus (AAV) capsids are used as gene delivery vectors in genetic therapies.^29^ AAVs naturally exist as multiple serotypes, each defined by differences in capsid protein sequences that determine tissue tropism and immunological responses, enabling their selection for tissue-specific treatments.^30,31^

Currently, solution-phase methods are the predominant commercial approaches employed for characterization of these complex, high-value analytes. The gold-standard technique, analytical ultracentrifugation (AUC), typically requires large sample volumes, suffers from low throughput, and demands a high degree of technical expertise.^32^ Faster and more sample-efficient methods such as anion-exchange chromatography (AEX) and size-exclusion chromatography coupled to multi-angle light-scattering (SEC-MALS), do not provide sufficient resolution to distinguish empty and payload-containing vectors from those carrying truncated genetic cargo. AEX performance in particular can also be serotype dependent.^33^ Another light scattering method, mass photometry (MP) makes use of the light scattered by particles attached to a glass surface.^34^ MP offers ease-of-use and speed (compared with e.g., AUC). The mass resolution of MP is low in comparison to MS approaches (R ∼ 3 (*m/*Δ*m*)),^35^ with a mass range of 0.03-6 MDa.^36^ A related technique, termed macro-MP, was developed to characterise larger analytes, although it does not directly provide mass information. Instead, macro-MP reports particle diameter and scattering contrast, which may be used as a qualitative proxy for low resolution mass comparisons if particles have the same diameter and composition.^36^

CDMS combines the sensitivity and resolution of traditional native MS with individual ion *m/z* and, crucially, charge (*z*) measurement, thereby overcoming the *m/z*-congestion problem in the mass analysis of large and heterogeneous analytes.^37^ In CDMS, the *m/z* and *z* are measured simultaneously by passing an ion through a charge-conducting cylinder (CC). When the ion enters the CC it induces a charge separation which is detected by a charge sensitive amplifier. For a CC with aspect ratio greater than 3, the induced charge is virtually independent of the ions trajectory through the CC and the charge can be measured accurately.^38–40^ For ions of known axial kinetic energy (KE), the time-of-flight through the CC is proportional to the square root of their *m/z* ratio. Thus, both the *m/z* ratio and *z* can be determined from a single measurement. This approach was first used in the early 1960s to measure the masses of microparticles.^39^ In the 1990s Fuerstenau and Benner adapted this approach to electrosprayed ions. These early experiments relied on passing ions through the CC just once, resulting in relatively low charge precision and limiting detection to ions carrying several hundred charges or more.^41,42^ Benner later positioned the CC between two electrostatic ion mirrors to enable ion trapping and multi-pass operation.^43^ A precision of ∼2 elementary charges (*e*) root mean square deviation (RMSD) was achieved. This electrostatic linear ion trap based CDMS (ELIT-CDMS) approach was later improved by Jarrold and co-workers, who introduced the use of fast Fourier Transforms (FFT) to analyse time domain data from the trapped ions.^44^ In this method, the *m/z* is determined from the oscillation frequency, and the charge is obtained from the FFT magnitudes. Subsequently, the charge limit of detection was reduced to 1 *e*^45^ and baseline resolution between ions that differ only by a single elementary charge (charge precision RMSD of ∼0.2 *e*) was demonstrated for ions trapped for ∼3 s.^46^ More recently, this level of performance was achieved within ∼0.95 s as a result of improvements in amplifier and trap design. Cryogenic cooling of the amplifier further reduced this to ∼0.65 s.^45,47,48^ Since charge is quantized, unit-charge resolution allows for near-perfect charge assignment for all detected ions. When the ion’s charge can be assigned with a low error rate, the precision in mass measurement is limited only by imprecision in *m/z* determination.^46^

Different iterations and implementations of CDMS have also been realised by Gamero-Castaño, Williams, and Dugourd *et al*.^49–51^ Notably, the use of conical ion mirrors and an array of CCs together with appropriate data-processing enabled CDMS measurements of ions having a wide spread of KEs. In this approach, harmonic ratios for individual ions are used to infer their KE, allowing the correct frequency to *m/z* relationship to be determined. These techniques are particularly enabling for experiments involving dissociation of ions inside the CDMS trap.^51,52^ Another CDMS implementation featuring dual sector design based on Poschenrieder’s geometry^53^ was proposed by Hoyes *et al.* and is understood to be currently undergoing development.^54^

To enable accurate charge measurement, it is essential to determine whether an ion changes charge state or disintegrates during the trapping event. In the ELIT-CDMS implementation of Jarrold and co-workers, this is achieved by performing a series of short-time, overlapping, fast Fourier transforms (ST-FFTs) which provide temporal information, allowing the application of “quality filters” that reject ions exhibiting charge loss, fragmentation or other instabilities.^44,55^ An alternative approach termed Selective Temporal Overview of Resonant Ions (STORI) has been demonstrated by Kafader *et al*.^56^ Here, sections of the time-domain data of increasing length are transformed with FFT, and the resulting magnitude is plotted against time. For ions which did not survive the entire trapping event, the resulting STORI plot exhibits two distinct regions: a linear region (slope ∝ charge), and a flat region corresponding to the time intervals during which the ion was no longer present. The STORI approach has been employed in the Direct Mass Technology^TM^ (DMT) mode which is available on selected Orbitrap^TM^-based instruments.^57^

The use of conventional ion-trapping *m/z* analysers with inductive detection (e.g., FT – Ion Cyclotron Resonance, FT-ICR) for charge determination of single-ions was first conceived by Smith *et al*.^58^ However, both FT-ICR and Orbitrap analysers present fundamental challenges for high-precision charge measurements. Unlike the CC used in CDMS, in these devices the detection electrode does not surround the ion, so the measured signal (∝ charge) depends on an ions’ trajectory. Moreover, because these analysers were not optimised for precise charge measurement, long transients (seconds to tens of seconds) are required. The best charge precision reported on Orbitrap-DMT platform is approximately 0.55 *e.* This performance was achieved by Heck and coworkers using an aftermarket data acquisition system after 24 s of ion trapping.^59^ In contrast, the most commonly used iteration of ELIT-CDMS from the Jarrold lab achieves 0.55 *e* in just 0.3 s, with more recent improvements reducing this to approximately 0.1 s.^46,48^ Extended transients exacerbate practical challenges, including KE loss (through ion-gas collisions), charge reduction, and changes in the mass of ions during a transient.^59,60^ These factors, together with the transmission profile of the front-end ion optics, limit the mass resolution and applicability of Orbitrap-DMT to analytes below several MDa, although recent developments aim to extend this range.^61^

While the primary motivation for precise *z* and *m/z* determination is improved mass precision and accuracy, these measurements can also provide valuable insight into the conformational properties of each analyte ion. In particular, extended particles are generally able to carry more charges than more compact conformers.^6,8^ The relationship between mass and charge has been previously studied by de la Mora, who demonstrated that for a given chemical class (e.g. proteins, synthetic polymers), deviations from the theoretical maximum charge (Rayleigh limit)^62^ can be used as an indicator of particle compactness or globularity.^63^ CDMS data can conveniently be represented as scatter plots of *m* and *z* which readily show differences in charging behaviour for particles of the same mass but different conformations.^64^ This is expected to be a powerful tool for structural characterisation of ions, providing conformational information analogous to that obtained from ion mobility experiments.^65^

Collectively, the above factors motivated the development of the first commercial mass spectrometer incorporating an ELIT-CDMS analyser. During the design process, emphasis was placed on useability, encompassing both hardware and software, to make high-resolution, high-mass, native MS measurements accessible to a wider scientific community, including researchers not traditionally trained in MS.

## Instrument Design and Operation

The ELIT-CDMS analyser was integrated into a newly-designed, compact, benchtop nESI-MS instrument (Figure 1). Briefly, ions are generated using a nano-electrospray (nESI) ion source and directed toward an inlet tube. Ions pass through source-transfer ion optics based on StepWave™ ion guide technology. RF-only segmented quadrupoles are then used to transmit ions into subsequent differentially pumped regions. Transfer and steering lenses then guide and accelerate the ions into the ELIT-CDMS analyser. Each of the above components is described in detail in the sections below.

**Figure 1.**
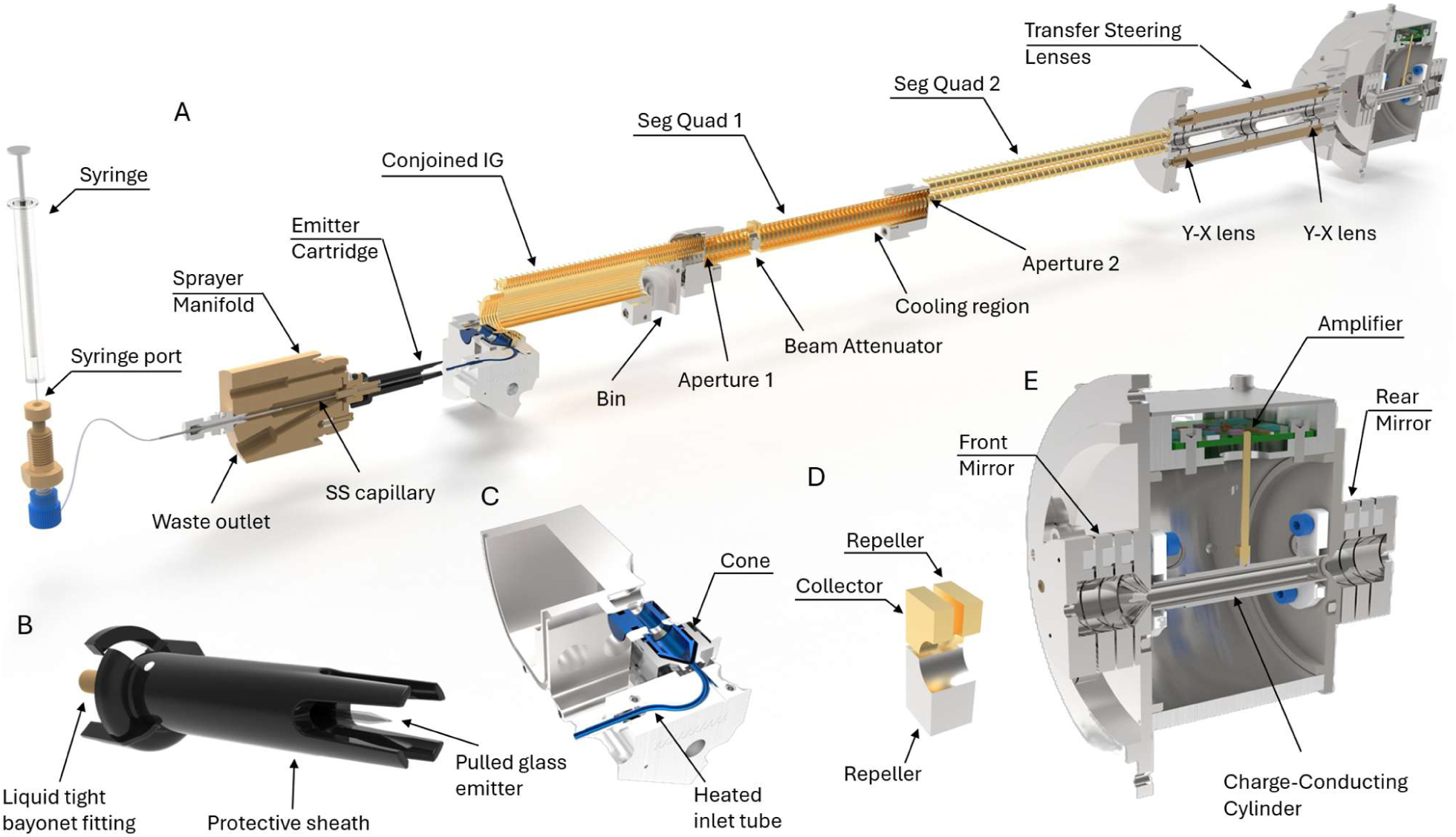
Instrument design: (A) instrument layout overview; (B) emitter cartridge; (C) ion inlet components; (D) beam attenuator electrodes; (E) electrostatic linear ion trap.

### Ion Source

Novel nESI emitter cartridges were developed to leverage the benefits of pulled glass emitter nESI technology, while addressing its key usability challenges. Emitter cartridges are made from an injection-moulded conductive polymer which houses and protects the glass needles, see Figure 1 and Figure S1A in Supporting Information (SI). This emitter sheath acts as a counter electrode for the spray together with the MS inlet tube, reducing the need for the positional adjustments. A liquid-tight bayonet fitting and electrical connection is located at the opposite end (Figure S1B in SI). Once installed onto a sprayer manifold, the internal (stainless steel, SS) capillary penetrates the glass emitter (Figure S1C in SI). The opposite end of this capillary is connected to the syringe port via a PEEKsil^TM^ section (Figure 1A). The SS section of the capillary delivers sample to the tip of the glass emitter and provides the electrical connection to the sample solution. This design allows multiple samples to be analysed using the same emitter, without the need for voltage and position adjustments for each sample. Typically, 10 µL of sample is introduced through the syringe port to fill the 5 µL capillary dead volume and the emitter tip. Between different samples, the emitter is flushed using a 250 µL syringe. Waste liquid flows through the opposite end of the emitter and into the waste outlet, which is connected to a waste bottle. Since the wash solutions used here contain high organic solvent content and are used in volumes far exceeding that of the sample, the resulting waste consists predominantly of wash solution, effectively inactivating biohazardous material. A sterilising agent may be added as an additional precaution. Very low volumes (<2 µL), common for extremely high-value samples, can be loaded offline using a streamlined version of the traditional nESI workflow (Figure S2 in SI).

As some of the analytes of interest may be biohazardous, the aerosol created by nESI must be contained. The enclosure features an active extraction system with an Ultra-Low Particulate Air (ULPA) filter and interlock switches (Figure S3 in SI). When the sprayer manifold is retracted (e.g., for emitter replacement, see Figure S4A in SI), high voltage is disabled and the extractor fan runs at high speed, sweeping the enclosure and exhausting it through the ULPA filter. This is similar to the operation of a biological safety cabinet. When the emitter is installed and the manifold re-inserted (Figure S4B in SI), the blower switches to low speed to maintain gentle evacuation without disturbing the nESI spray.

### Ion Optics

Ions are introduced into the vacuum system through an inlet tube embedded in a heated aluminium block (Figure 1C; <300 °C). Following the inlet tube, ions pass through the inlet cone before being redirected into an orthogonally oriented conjoined ion guide (IG)^66^ operating at ∼6 mbar. The lower portion of the conjoined IG steers the charged particles towards the upper, stacked ring portion (330 kHz, <400 V_pp_). Ions are subsequently guided through the first differential aperture (Aperture 1 in Figure 1A). The following ion guide is a segmented quadrupole (Seg Quad 1 in Figure 1A; 270 kHz, <700 V_pp_, ∼0.15 mbar) with DC gradients applied to the first section and the following cooling region.

This guide also contains a beam attenuation device, see Figure 1D. It comprises three electrodes which can be switched between two states (Figure S5A in SI). In an ON state the continuous beam of charged particles is transmitted through. In the OFF state, the beam is deflected and hits one of the electrodes. The duty cycle of switching between these states determines the fraction of ions progressing towards the subsequent sections of the instrument. The “train” of ion packets leaving the beam attenuator is slowed down in the cooling region (Figure 1A) allowing for re-merging and formation of a continuous ion beam (Figure S5B in SI). This design enables deterministic control of the ion transmission.^67^ The ions then pass through the second differential aperture (Aperture 2 in Figure 1A) and the subsequent, segmented quadrupole (Seg Quad 2 in Figure 1A; 310 kHz, <600 V_pp_, ∼5e-5 mbar) towards acceleration and steering optics.

The ELIT device has been designed to accept ions with axial kinetic energy (KE) of 130 eV/z.^58^ The width of the KE distribution is an important factor in determining the available *m/z* resolution (see Figure S6A in SI). Therefore, it is beneficial to accelerate ions at high vacuum, to minimize dependence of resultant ion KE on collision cross-section. Here, we accelerate ions through a stack of DC-only elements (Transfer Steering Lenses, Figure 1A) operating at ∼5e-7 mbar. This region features two sets of focusing and X-Y steering lenses which are designed to correct for any parallel and axial mechanical misalignment between the ELIT trap and the upstream ion optics.

### Electrostatic Linear Ion Trap

The design of the ELIT device (see Figure 1E and Figure 2A) is based on that developed by the Jarrold group.^58^ It is co-axial with the nominal ion path and features a charge-conducting cylinder (CC) located between two ion mirrors. The front and rear ion mirrors (FM and RM) are each comprised of three ring electrodes, and enable stable trapping trajectories for ions passing through the CC. The trapped ions’ oscillation frequencies can change during the trapping event owing to inevitable collisions with the background gas (ELIT operates at ∼3e-9 mbar). In addition, individual ions will take slightly different trajectories through the trap as a result of their initial phase space distribution. This geometry was chosen to allow axial transmission while also fully encapsulating a charged particle so that the measured image charge accurately reflects the charge on the ion. This design effectively removes trajectory and KE dependences on the charge measurement, which is consequently limited only by electronic noise. Multiple ions may be measured simultaneously provided that they have sufficiently different *m/z* values (i.e., frequencies). Additional data processing approaches may be employed to recover signals which overlap in the frequency domain.^69^

**Figure 2.**
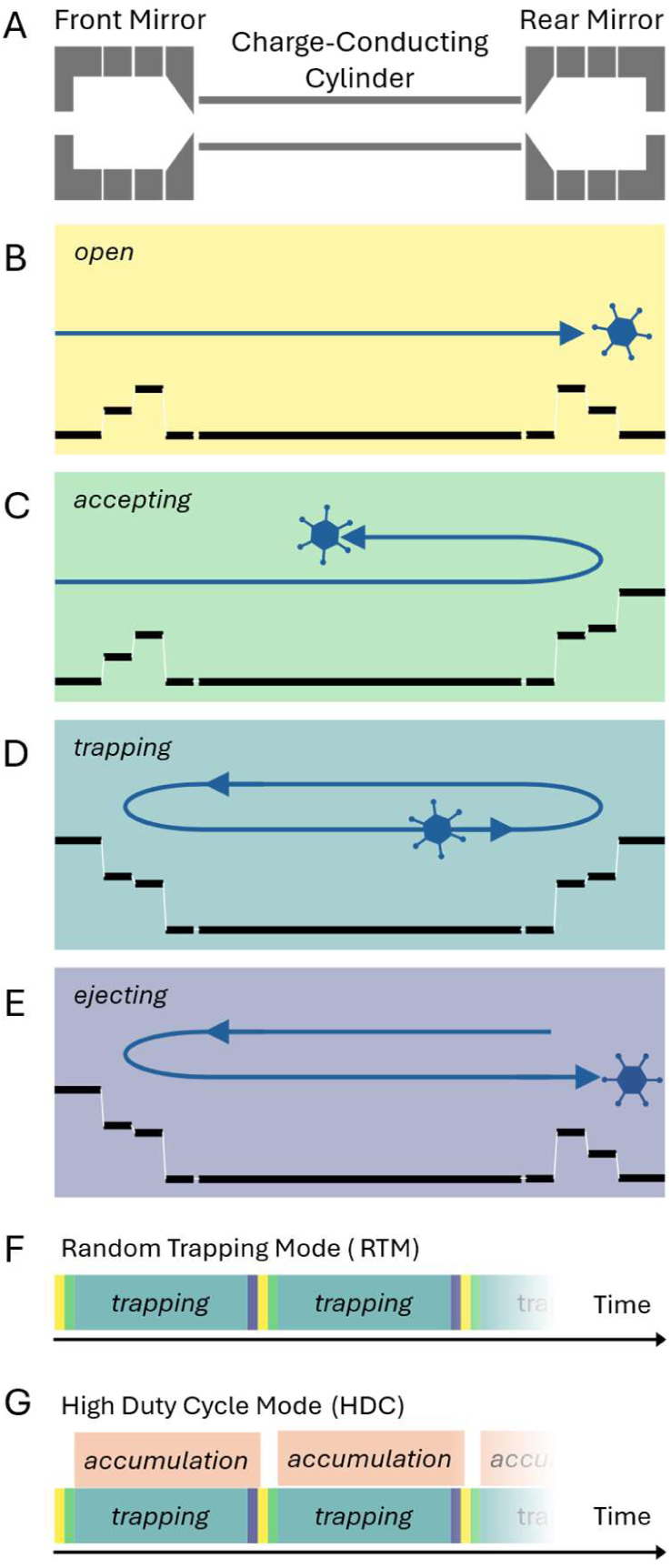
ELIT operational states and acquisition modes: (A) schematic of ELIT elements; (B-E) ELIT operational states; (F, G) experimental sequences showing duration of different operational states in RTM and HDC acquisition modes, respectively.

Ion trapping stability and *m/z* resolution are dependent on the phase space of the ions entering the ELIT (i.e., radial offset and angular divergence, see Figure S6 in SI). Great care was taken to design a trap that enables precise measurements of *m/z* while providing a degree of energy and trajectory independence.^68^ The lengths and voltages applied to the FM, CC and RM were optimised such that each trapped ion spends 50% of its time inside the CC and the remaining 50% outside of it (i.e., within the FM and RM). The resulting, single-ion signal closely resembles a square-wave. This symmetric signal has a simple Fourier series representation – the energy is distributed exclusively across the odd harmonics. Since the ion energy is known and fixed, the oscillation frequency (*f*) of a single ion signal can be related to its *m/z* ratio via Equation (1) below:^44^

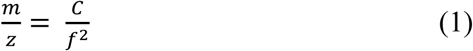

Where the geometry and voltage dependent constant *C* is determined during the calibration process.

Operation of the ELIT is realised through sequential switching between 4 states (*open*, *accepting*, *trapping* and *ejecting*). This is achieved by changing voltages applied to the FM and RM, as illustrated in Figure 2B-E. Ions can enter the trap when the FM is OFF in both the *open* and *accepting* states (applied for 5 and 1 ms, respectively). When the ELIT switches to the *trapping* state, all ions present within the CC and RM at that moment become trapped and subsequently analysed. Because ions are captured in a stochastic manner under these conditions, this mode of operation is referred to as Random Trapping Mode (RTM). The duration of the *trapping* state is typically set to 0.1 to 2 s. Following trapping, ELIT voltages switch to the *ejecting* state (1 ms) and the ions are released from the ELIT through the RM. This sequence is repeated throughout the acquisition time as shown in Figure 2F.

When coupled with a continuous ion source such as ESI, the ion utilization rate (i.e., duty cycle) is defined as the ratio of residence time within the trappable region of ELIT to the total sequence cycle time (t_total_). The residence time may be expressed in terms of the oscillation period *T*. Since ions may be reflected within RM prior to being trapped (Figure 2C), the effective trappable length constitutes the length of FM + CC + RM + RM + CC. This corresponds to 1.75 oscillation periods *T*. Note that only a single FM length was included – if an ion happens to be inside the FM when it switches state (*accepting* → *trapping*), the resulting change in the electric field will alter its KE, affecting stability within the ELIT and the pattern of harmonics – these ions will be either lost or removed during data processing steps (as described later). Thus, the duty cycle may be calculated using Equation (2); the corresponding plot is shown in Figure S7 in SI.

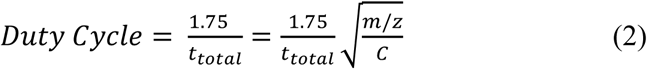

To improve the duty cycle, ions can be accumulated upstream of the ELIT for the duration of *open*, *trapping* and *ejecting* states and released just before the ELIT switches to the *accepting* state.^70^ Here, we accumulate ions in the cooling region of Seg Quad 1, by raising the potential applied to Aperture 2 (Figure 1A). Since the transit time between the cooling region and the ELIT is *m/z* dependent, the timing of ion release from the cooling region and opening of the ELIT trap determines the *m/z* range over which duty cycle enhancement is achieved. This method is expected to achieve near 100% duty cycle for a narrow *m/z* range (further details can be found in SI). To allow enhancement over a broader *m/z* range, a series of discrete, varying delays can be applied sequentially. This is referred to as High Duty Cycle (HDC) acquisition mode (Figure 2G).

### Signal processing and data acquisition

The charge induced by ions on the CC is superimposed on electronic noise (Figure 3A). The CC is directly coupled to a charge-sensitive amplifier (Figure 1E); its architecture originates from a high-frequency device developed by Bertuccio^71^ and was adapted to operate in the frequency range of interest (3-300 kHz).^48,72^ The key aspect enabling the ultra-low noise measurements presented here is the removal of the feedback resistor – a source of thermal noise. This amplifier circuit makes use of a JFET for its extremely high input impedance across the kHz bandwidth. The JFET, can self-bias itself to near 0 V but remain within a transconductance region that allows for amplification.

**Figure 3.**
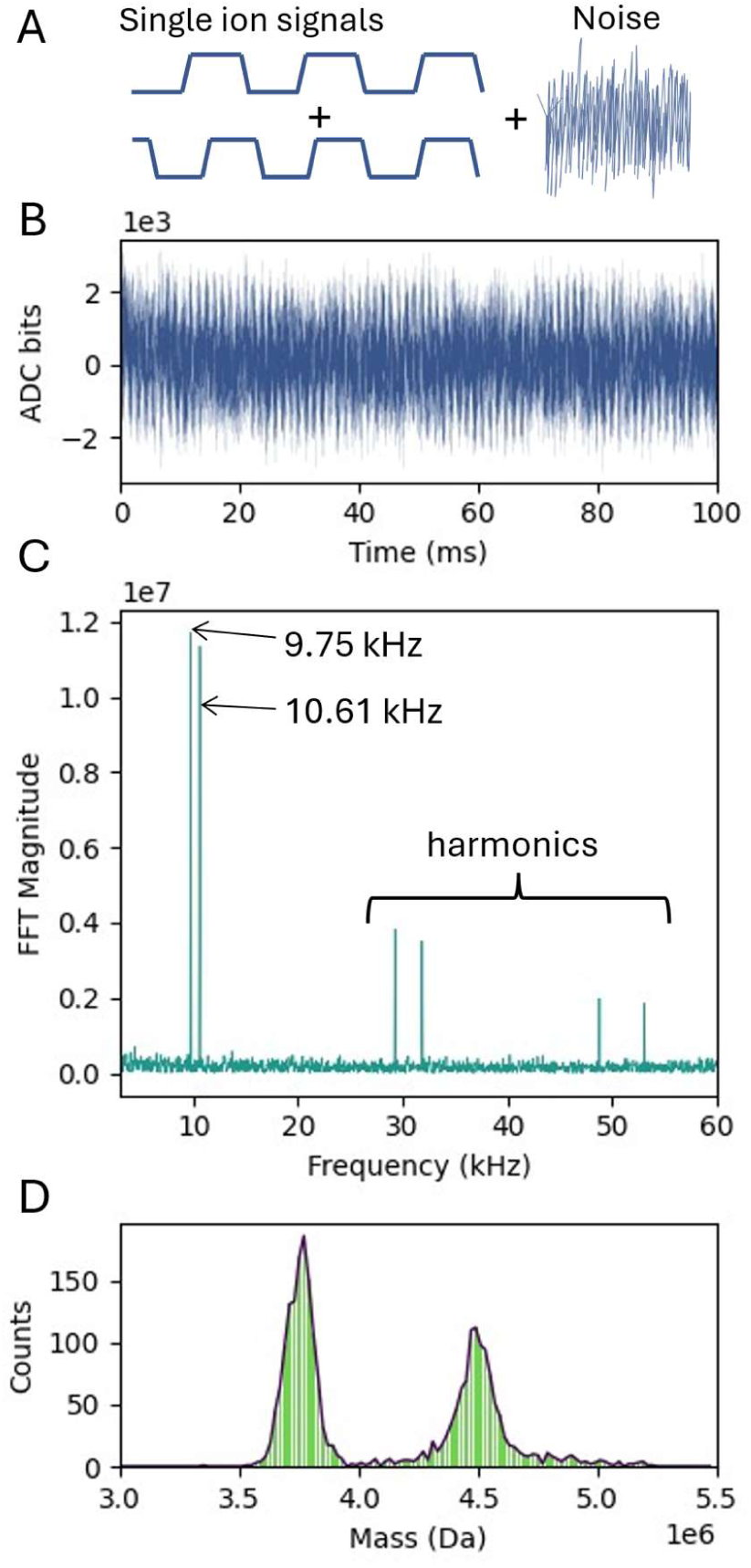
Data acquisition and processing workflow. A: Schematic illustration of the signal ion signals superimposed on electronic noise. B: Experimental time domain data. C: Data in frequency domain, following FFT. Two ion signals are observed. D: Mass histogram of *∼*2800 single ion measurements.

The general data acquisition approach used here (Figure 3B-D) follows that described by Draper and Jarrold and will be discussed only briefly here.^55^ The amplified signal is recorded over the trapping time (0.1 – 2 s) using a 2.5 MHz analogue-to-digital converter (Figure 3B). A short portion of the data (∼1 ms) at the start of the transient is discarded to remove pickup from FM switching. All FFTs are zero padded to a total of 2^18^ points and apodized using a Gaussian window. A “survey FFT” is performed on the first 2^16^ points (∼26 ms) of the time-domain data. Peak detection of the resulting frequency spectrum (Figure 3C) is performed to identify candidate ion signals. These candidates are then tracked through a series of shorter, overlapping FFT windows typically of length 2^15^ points (∼13 ms) offset by ∼1.3 ms to allow for changes in the ion population over time to be monitored. Statistics are collected for a variety of ion properties over these windows (including frequency measurements and charge standard deviation). Since ions may be lost during a transient, the survival time of the ion is also recorded. A linear fit to the frequency measurements as a function of time is performed, and an extrapolated value at time zero is used to calculate the measured ion frequency and, consequently, *m/z* ratio.

Charge determination is aided by a dynamic calibration approach, described by Todd and Jarrold.^47^ A small antenna located inside the ELIT enclosure generates a sine-wave of 250 kHz. The amplitude of this “internal standard” signal is monitored throughout the trapping event, as discussed above. An average charge is calculated for each ion by normalizing its amplitude to the internal standard signal and applying a linear calibration with an offset, determined during the charge calibration procedure (see SI). The remaining statistics are used to qualify the detected ions, and ions that do not meet these criteria are filtered out.

In the example dataset (Figure 3C), two signals are observed at 9.75 kHz and 10.61 kHz, with the peaks corresponding to odd harmonics present at higher frequencies. The method described above yields *m/z* values of 29,211 and 24,670 with the corresponding charge values of 157.1 and 149.1. Their corresponding masses are calculated as 4.589 MDa and 3.679 MDa. This process is repeated for all ions detected in each trapping event, and throughout the acquisition. Finally, all the obtained mass values are binned into a histogram, resulting in a mass spectrum (Figure 3D). All the above steps are performed in real-time, allowing for data (e.g., mass spectrum) updates every 1 s. Note that there is no deconvolution, smoothing or processing of the *m/z* or *z* values before multiplication. Any such processing would make the measurements “ensemble” rather than single ion.

### Mass precision and accuracy

Since precision in mass determination is limited by imprecision in the two primary measurements (*m/z* and *z*), it is of interest to evaluate their relative importance. Standard error propagation results in the following Equation (3), relating mass imprecision to the corresponding imprecisions in the *m/z* and charge measurements.

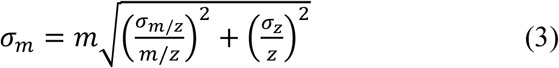

The equation above can be rearranged to highlight the relationship between *m/z-* and *mass-*domain resolutions. Figure 4 illustrates the corresponding trends, calculated for a range of charge imprecision values. The key insight from Equation (3) and Figure 4 is that high *m/z* resolutions are largely irrelevant unless charge measurement performance is sufficient to enable charge quantization. In the latter case, the charge-dependent term in Equation (3) vanishes, and *mass* resolution equals *m/z* resolution.

**Figure 4.**
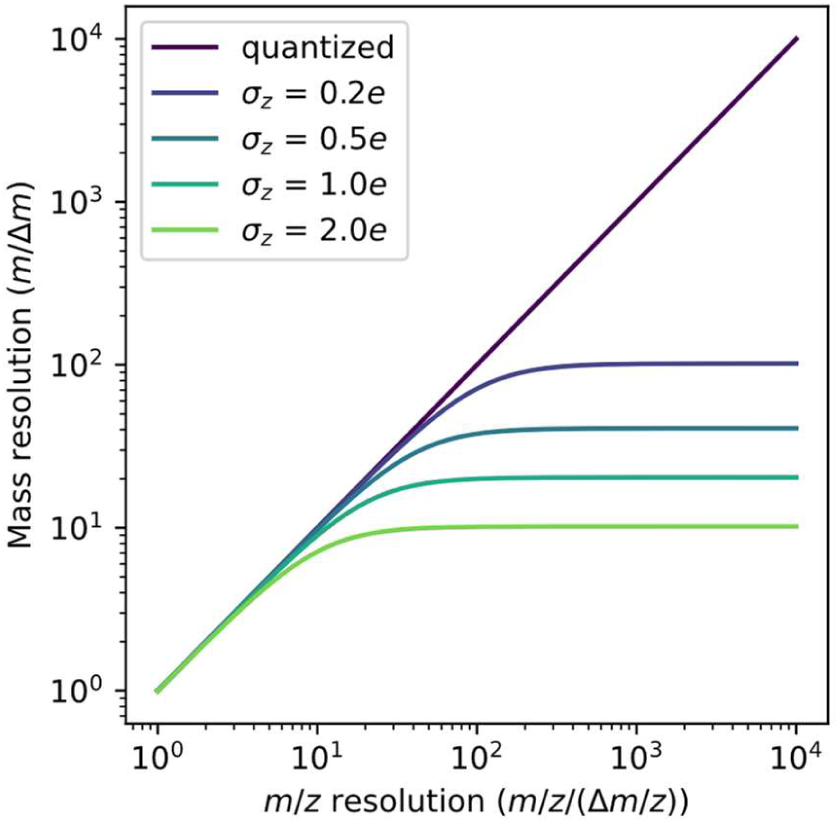
Relationship between m/z and mass resolution in CDMS analysers, according to Equation (3), plotted for an ion with 48 charges.

The imprecision in the charge measurement is reduced by increasing the measurement time according to:

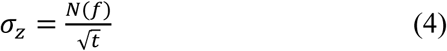

Where *N*(*f*) is related to the electrical noise at the ion oscillation frequency (*f*) and *t* is the trapping time. The frequency dependence of *N*(*f*) is similar to 1/*f*, so the noise is worse at lower frequencies. Equation (5) allows an estimation of the trapping time required to reach the quantization threshold (i.e., *σ_z_* = 0.2 *e*) for ions with a given oscillation frequency (i.e., *m/z*) using a reference measurement of charge imprecision (*σ_ref_*) for ions with the same oscillation frequency after a trapping time *t_ref_*.

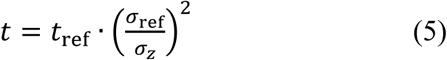

Similarly to precision, the accuracy of mass measurement using ELIT-CDMS is limited by accuracies in *m/z* and charge.^47,73^ Both these measurements require calibration, however, owing to their size, CDMS calibrants present unique challenges, and we have developed a new mathematical approach that is designed to address them rigorously.^73^ This will be detailed in a subsequent publication but will be briefly introduced below.

CDMS calibration relates the frequency and amplitude of the signal produced by individual ions to their *m/z* ratio and charge state, respectively. For each potential standard, a two-dimensional region (frequency and amplitude) of data is selected to isolate the charge states to be used for calibration. Individual peaks are located in this space using a specially adapted deconvolution algorithm. A mathematical model^73^ of the data has been constructed relating the theoretical mass to the detected peak positions. A probabilistic algorithm is then used to determine the resulting *m/z* and charge calibration parameters together with their uncertainties. Following the calibration procedure, the measured mass accuracy is expected to be below 1 %.^73^ Further details and example calibrant datasets are included in SI.

## Materials and Methods

All protein standards were purchased from Sigma-Aldrich. Dengue and chikungunya virus-like particles (VLPs) were obtained from the Native Antigen Company. AAV empty and full (with CMV promoter-driven expression of GFP) capsid standards of various serotypes (AAV6, AAV8, AAV9 and AAVrh10) were obtained from multiple commercial suppliers including Virovek, Charles River Laboratories, and AAVNergene. Supplier-provided viral titres ranged from 1 × 10^11^ to 2 × 10^13^ vp/mL for empty capsids and from 1 × 10^11^ to 2 × 10^13^ vg/mL for full capsids. 20-50 µL of each sample was directly buffer exchanged into 200 mM aqueous ammonium acetate (AmAc) solution (Invitrogen, AM9070G), pH 6.8. For AAVs and the chikungunya VLPs, 0.01% poloxamer 188 (P-188) non-ionic surfactant (Gibco^TM^, p/n 24040032) was added to the buffer exchange solution. Samples were prepared using either Micro Bio-Spin^TM^ P-6 size-exclusion gel columns (Bio-Rad^TM^, p/n 7326221) or Amicon^TM^ 100 kDa molecular-weight cut-off, ultra-centrifugal filters (Millipore^TM^, p/n UFC5100).

A development prototype of the Xevo^TM^ CDMS instrument was used for all experiments. 10 uL of each sample was aspirated by a 25 uL gas-tight syringe (Hamilton^TM^, p/n 80265) and injected into the emitter cartridge via the syringe port of the nESI source. Wash solutions were (A) deionized water, (B) magic mix (1:1:1:1 mixture of water : isoproponal : methanol : acetonitrile), and (C) 200 mM AmAc + 0.01% P-188. The wash sequence (A, B, A, C) was performed between each sample injection, using 250 uL syringe (Hamilton, p/n 81165). Ions were generated in positive polarity nESI, with spray voltages ranging from 1.3 to 1.7 kV and subsequently trapped for 0.1 s, unless otherwise stated. Data acquisition times for the different analytes were typically between 5 and 20 minutes (stated in figure captions). Screenshots of the instrument control software and calibration page are shown in Figure S8 in SI. Signal processing and visualization were performed using waters_connect™ CDMS Toolkit Software. Custom data plots and figures used in this manuscript were generated using Python^TM^ plotting libraries.^74,75^

## Results

### Charge precision enables high-resolution *mass* measurements

Figure 5 shows representative CDMS data obtained for β-galactosidase ions trapped for 0.1 and 2 seconds. The panels in Figure 5 show the *m/z* histogram (top left), a scatter plot of *m/z* and charge (bottom left), a mass histogram (top right) and a charge histogram (bottom right). After trapping for 0.1 seconds (light green data), the *m/z* resolution (*m/z*/Δ(*m/z*)), measured on the most intense peak of the CSD, is ∼160. The charge RMSD, measured on the same (*m/z* resolved) peak is ∼0.93 *e*. The mass resolution (*m*/Δ*m*), measured on the mass histogram (top-right panel) is ∼20. Trapping ions for 2 seconds (dark green data) has no effect on *m/z* resolution – as expected for single-ion ELIT-CDMS measurements. According to Equation (5), a trapping time of ∼2 s is required to reach the charge quantization threshold in this instance. Experimental data corroborates this prediction: after 2 s of trapping, the average charge RMSD across the charge state distribution is 0.20 *±* 0.03 *e* (see Figure S9A in SI). Mass resolution is now increased 4x (∼80), in line with Equation (3) (see Figure S9B in SI). To “quantize” the charge, ions within ±0.15 of the midpoints between integer charge state values were discarded, and the remaining charge state values were rounded to the nearest integer (quantized, purple data).^46^ This further increases the mass resolution to ∼160, matching the *m/z* resolution (as discussed previously, Figure 4). As can be seen from the scatter plot, there are few instances where quantization resulted in the wrong number of charges being assigned, these correspond to ∼1 % of the total number of ions.

**Figure 5.**
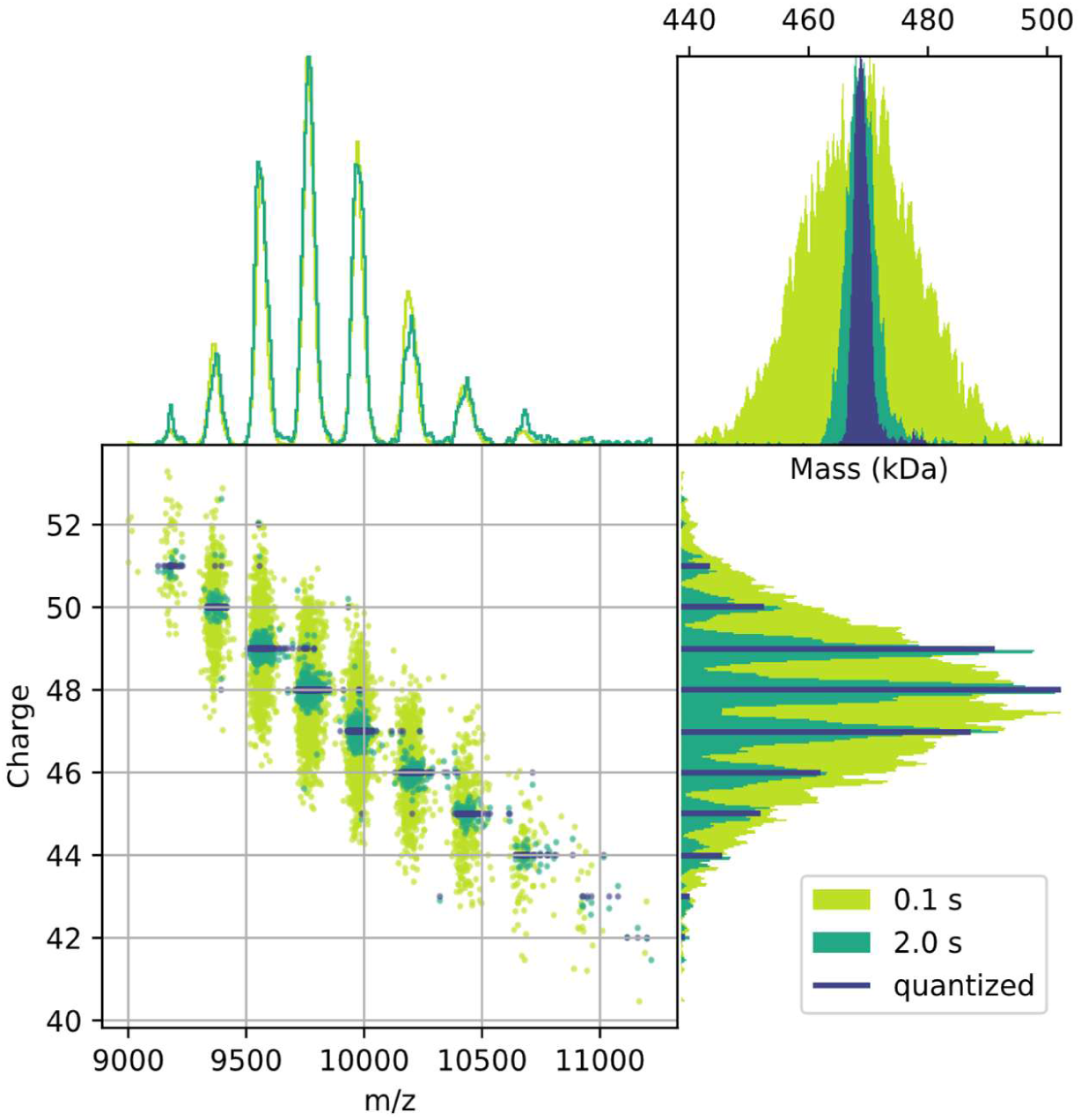
Representative CDMS data highlighting instrument performance following 0.1 s and 2 s trapping (light and dark green series, respectively) demonstrated on β-galactosidase ions. The purple series corresponds to 2 s data after charge quantization (details in the main body). Top left: m/z spectrum. Bottom left: scatter plot of m/z versus charge, each point represents a single ion measurement. Top right: mass distributions*. Bottom* right: charge distributions. Data at 0.1 s and 2 s of trapping was acquired for 30 and 240 minutes, respectively.

### Mass range, *m/z* transmission and trapping stability

An important requirement of the instrument presented here is the ability to transmit ions spanning a broad *m/z* range. Figure 6 shows data obtained from a sample containing Chikungunya (CHIKV) VLPs. This VLP is used to mimic the structure of the CHIKV virus, an alphavirus with a T=4 icosahedral symmetry. CHIKV VLP consists of 240 capsid proteins and ∼480 envelope proteins (240 × E1 and 240 × E2), with estimated total protein mass of ∼30 MDa. CHIKV VLP also include a host-derived lipid bilayer (phospholipids, cholesterol, sphingolipids) with estimated total mass of 10-15 MDa. Furthermore, E1 and E2 proteins are heavily glycosylated, with estimated total glycan mass of 3-5 MDa. Therefore, the estimated total CHIKV VLP mass is 45-55 MDa. The experimental CHIKV VLP data were plotted as *m/z* vs charge (Figure 6A), to highlight operational limits (i.e., *m/z* and *z*). As can be seen, the *m/z* distribution alone is very crowded and provides little insight into the nature of the species present. When plotted as mass vs charge, a series of distinct features are observed in the mass domain (Figure 6B). The most abundant peak at 52.3 MDa is attributed to the intact, monomeric CHIKV VLP. A series of additional smaller species are observed below 25 MDa which we attribute to trapped intermediates, degraded capsid assemblies and capsomers. Interestingly, larger species are also observed at approximately 104, 156, 208, 260 MDa. These species are absent in freshly prepared samples (“CHIKV VLP fresh”, see interactive Figures A and B and follow the instructions provided in the SI), suggesting that they exist in the solution phase and are not formed during the ESI process. The regular increase in mass suggests that these species correspond to aggregates. Each of the observed species spans a very broad charge (and *m/z*) range. For example, monomeric CHIKV VLP species are observed between *m/z* 80,000 and *m/z* 200,000, corresponding to the number of charges between 250–700. The upper charge measured for the monomeric and <25 MDa species broadly follows the theoretical prediction of the Rayleigh limit (Figure 6A, B, blue traces).^62,63^ In contrast, larger oligomers appear to exceed these predictions. This deviation may be attributed to the shape anisotropy (i.e. deviation from spherical geometry) and irregular surface topology, both of which may result in the accommodation of a greater number of charges. Overall, this dataset demonstrates transmission of ions over a mass range of up to 350 MDa, an *m/z* range of 6,000–250,000 and a charge range of up to ∼2000 charges. Data on smaller proteins (myoglobin and enolase) are included in Interactive Figure B (SI) and demonstrate transmission at *m/z* <6000.

**Figure 6.**
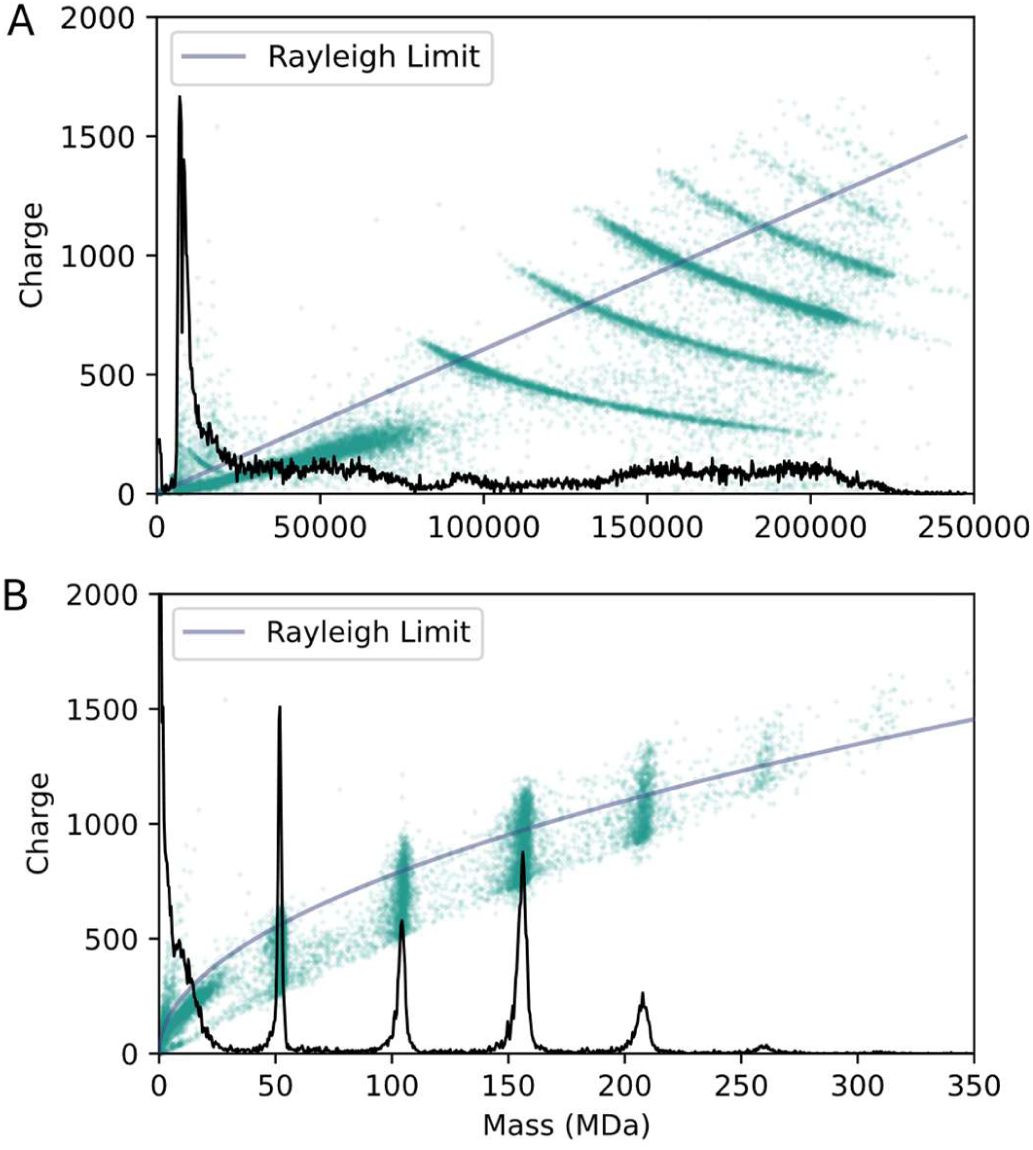
Mass and m/z transmission profile as observed on Chikungunya VLP and its aggregates. A: m/z spectrum and corresponding scatter of m/z vs z. B: Mass spectrum and corresponding scatter of mass vs m/z. Data was acquired for 80 min.

Prolonged trapping inevitably leads to ion-gas collisions and losses in ion’s KE. Since the stability of ions inside the ELIT analyser is energy dependent (Figure S6D) prolonged trapping may lead to losses. To verify this behaviour, duty-cycle corrected ion survival rate was plotted against trapping time (see Figure 7). For β-galactosidase ions, approximately 30 % of ions survived trapping for 2 seconds. Larger ions are more stable as evidenced for AAV particles; approximately 70 % of AAV ions are retained after 2 seconds. This is expected, since the loss of KE following collision (i.e., momentum transfer) will be smaller for heavier molecules. Most importantly, minimal differences in the survival of empty and full particles are observed. This contrasts with the earlier work of Ebberink *et al*., who reported that survival differences led to changes in apparent empty/full AAV ratios during extended transient experiments (1–24 s).^76^ In the present study, the higher performance of the charge detection system enables substantially shorter trapping times (0.1–2 s), thereby minimising such effects.

**Figure 7.**
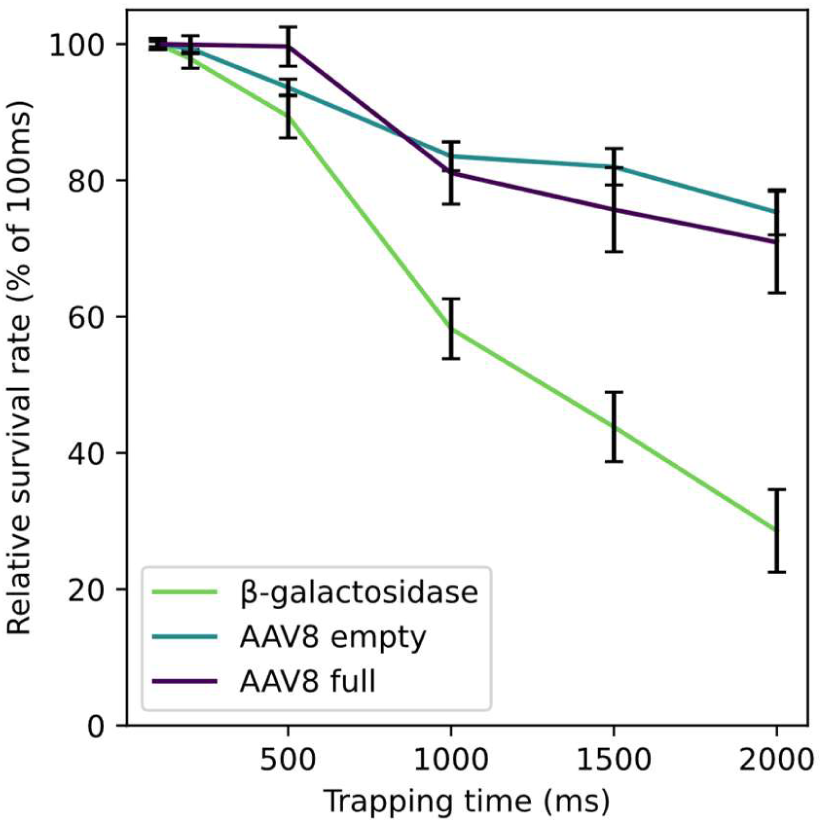
Stability of ions inside ELIT as a function of trapping time. Relative survival rate is defined as the duty-cycle–corrected ion arrival rate, expressed as a percentage of the ion arrival rate measured at a 100 ms trapping time. Ions with mass ranges of 0.44-0.50, 3.5-4.0 and 4.3-4.7 MDa were used for the corresponding β-galactosidase, AAV8 empty and AAV8 full traces. Error bars correspond to standard error.

### Analysis of adeno-associated virus capsids (AAVs)

Accurate relative quantification of empty and full AAV capsid content represents a critical quality attribute in AAV characterisation. Empty capsids, generated as product- and process-related impurities during vector manufacturing, can influence both immunogenicity and therapeutic efficacy. Reported levels of empty capsids range from 20% to over 90% in some preparations, contributing to batch-to-batch variability, elevated immune responses, and reduced transduction efficiency. Consequently, serotype-agnostic, precise analytical profiling of AAV capsid subpopulations is essential for the development, control, and comparability assessment of AAV-based therapeutics. In this section, we assess the performance characteristics of the instrument relevant to AAV analyses.

Reusing a single emitter cartridge enables reproducible analysis of multiple AAV samples. Cartridges do have a finite lifetime and need to be replaced when clogged by particulate material or damaged (e.g. by excessive voltage), it is relatively common to perform up to 10 injections of an AAV standards using the same cartridge. Figure 8 shows mass spectra recorded from ten injections (10 µL each) of a sample containing empty (E), full (F) and overfull (F*) AAV8 particles (from Virovek). Samples and flushes were injected using syringes (Materials and Methods). The sprayer position and applied voltage (1.4 kV) were kept constant. The appearance of the spectra remained consistent with all three capsids forms detected (E: 3.6-4.3 MDa, F: 4.5-5.0 MDa, F*: 5.0-5.8 MDa); the variation of the total (absolute) peak area was approximately 10 % across all injections. The standard deviation in relative peak areas was below ∼1 %, indicating a high level of precision in E/F/F* ratio quantitation even at relatively short acquisition time (5 minutes).

**Figure 8.**
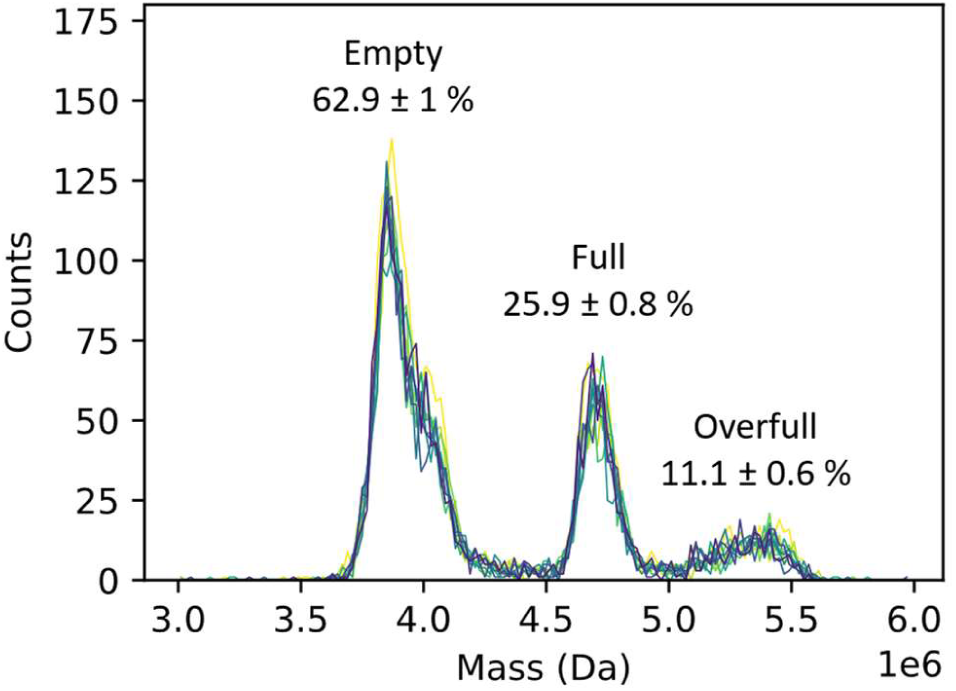
Multi-use emitters yield consistent spectrum quality for AAVs. Overlaid mass spectra from ten injections of AAV8 empty-full mixture (5×1012 vp/mL, from Virovek), using the same emitter cartridge. The intensity axis has not been normalized to highlight the variation in absolute abundance across injections. Labels show the relative mean peak areas (%) as well as standard deviations from 10 injections. Each sample injection was acquired for 5 minutes with beam attenuation set to 95%.

The sample concentration of ∼5 × 10^12^ vp/mL (as used for the AAVs shown in Figure 8) is near the limit of concentration for AAV formulations; further increases in concentration led to a high proportion of ions failing the quality filter. To investigate the lower concentration limit, the above sample was diluted 5- and 100-fold. This enabled asessment of the reliability of ratiometric measurements across the broader concentration range as well as comparison between RTM and HDC acquisition mode (see Figure S10A in SI). In RTM, using sample concentration of 1 × 10^10^ vp/mL (no beam attenuation), approximately 2000 ions were collected in 170 minutes within the AAV mass range (3-6 MDa). With HDC mode enabled (range set to *m/z* 20,000-40,000), the same number of ions was collected in ∼8 minutes. Collectively, these results demonstrate an approximate 20-fold enhancement in sensitivity between RTM and HDC mode across this *m/z* range. Consistent with the observations above, beam attenuation did not result in significant differences in the relative abundance of E, F, and F* species (see Figure S10B in SI).

Relative quantitation of the E/F ratio was further investigated across a range of E-F mixtures, prepared from another set of stock solutions (from Charles River Laboratories), containing predominately E and F species (Figure 9). The data were plotted as relative peak area versus the nominal E/F ratio (i.e., solution concentration), for each mixture. There is a good agreement between the E/F ratio in the prepared mixtures and the percentage of E and F capsids measured by CDMS, with a linear response and strong correlation observed (R^2^> 0.98) for the primary species.

**Figure 9.**
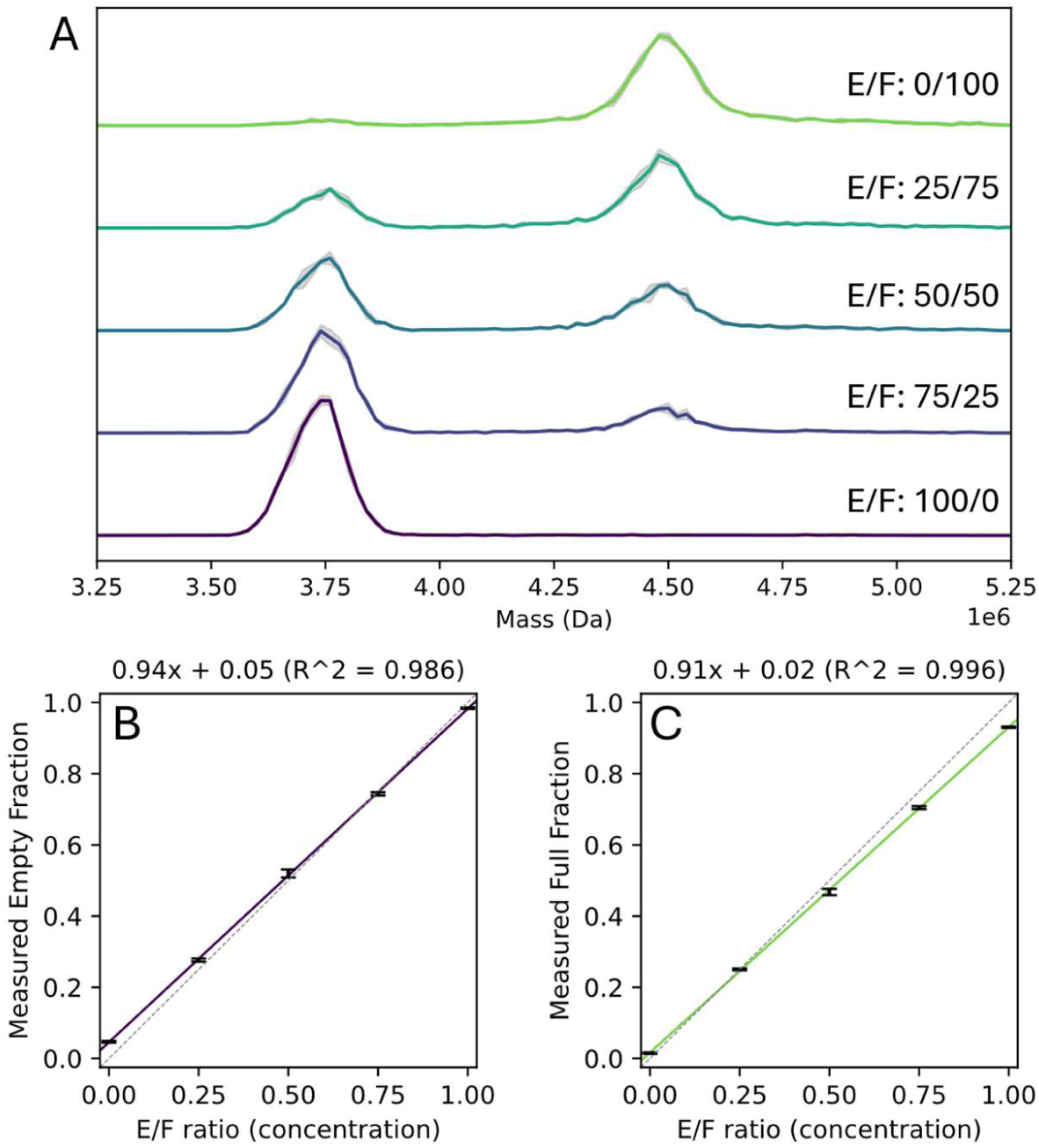
AAV8 ratio quantitation. A: CDMS spectra of E/F mixtures (1.5×1011 vp/mL each, from Charles River Laboratories). B: Measured area of peak corresponding to empty capsids (3.5-4MDa) vs nominal E/F ratio. C: Measured area of peak corresponding to full capsids (4-5MDa) vs nominal E/F ratio. Equations and colored solid lines in B and C correspond to linear fits to the experimental data (points). Grey area reflects standard deviation in each mass bin, from three replicates. Grey dashed lines in B and C correspond to 1-1 correlation. Each sample injection was acquired for 10 minutes.

Different AAV serotypes can exhibit distinct behaviour during capsid assembly, genome packaging, and purification as well as during the purification process. These variations can result in differences in capsid stability and E/F ratios, which directly impact the measured mass, charge distributions, and overall heterogeneity observed. Data obtained for several AAV serotypes (from AAVNergene) are presented in Figure 10. While minor differences in the masses of E and F species are expected across the serotypes, a markedly distinct patterns are observed for partially-filled (P) capsids (4–5 MDa). Here, performance of the instrument enables the resolution and identification of different subpopulations of P particles within each serotype; these may be of bioprocess, biological or clinical (i.e., immunogenic) importance.

**Figure 10.**
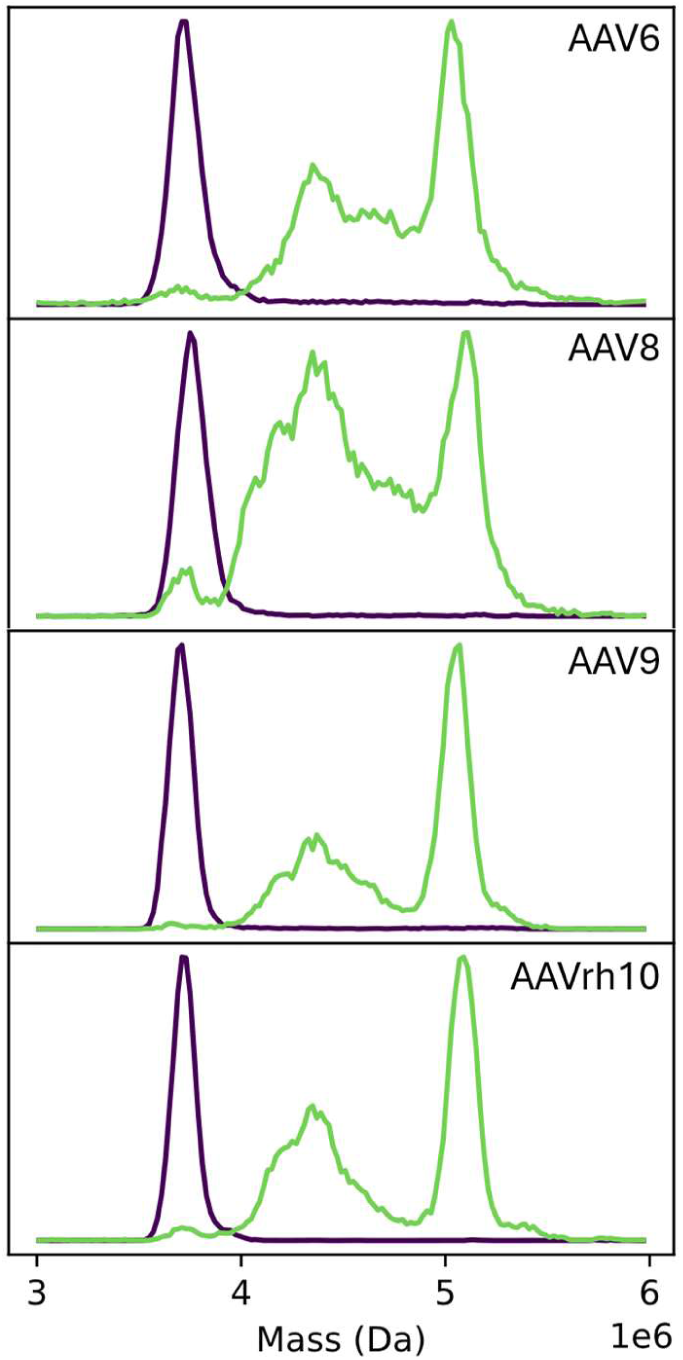
CDMS analysis of a selection of AAV serotypes from AAVNergene. Purple traces correspond to samples containing empty capsids, while green traces represent samples containing full capsids; the presence of empty and partially filled species in these samples reflects residual impurities. Each sample injection was acquired for 10 minutes.

### Inlet heating induces envelope loss and dissociation of virus like particles (VLPs)

Dengue virus (DENV) is a small, spherical, enveloped, flavivirus with a single-stranded RNA genome and icosahedral-like symmetry. It is the causative agent of dengue fever, a mosquito-borne disease that is prevalent in sub-Saharan Africa and other tropical and subtropical regions. The native virion consists of a lipid bilayer embedded with 180 copies of the envelope protein (En, 53.9 kDa) and the precursor membrane N-linked glycosylated protein (prM, unglycosylated mass 18.4 kDa).^77^ These components assemble within a host-derived lipid membrane to form particles consistent with T = 3 icosahedral symmetries. Based on this stoichiometry, the expected mass of a fully assembled VLP is ∼13 MDa (∼50 nm).^78^ DENV VLPs are known to be smaller (29-34 nm), non-infectious assemblies that lack the viral RNA genome, thereby eliminating the ability to replicate or cause infection.^79^ As a result, these particles are widely used for studying immune responses, supporting vaccine development, and serving as diagnostic reagents for the detection of dengue-specific antibodies.^80^

Here, we explored the applicability of the heated inlet for investigating thermal activation of the DENV VLPs. At room temperature, DENV VLPs are detected predominately at 7.1 MDa, suggesting that this preparation contains mainly “small” VLPs, instead of the 180-copy architecture of the wild-type viron (see Figure 11A). Several other species are observed at lower masses. Two sharper features are detected at 82 kDa and 172 kDa (see Figure 11B). These masses are close to the expected masses of prM-En heterodimer and its dimer, [prM-En]_2_. Broader, lower intensity features at 260 kDa, 340 kDa indicate presence of [prM-En]_3_ and [prM-En]_4_, respectively. Low mass features are convolved with elevated background signal between 40 kDa and 600 kDa, which may be attributed to constituents of viral envelope. Upon heating the inlet tube to 180 °C, minor decrease in FWHM is observed, and can be attributed to more efficient desolvation. At inlet temperatures between 180 °C and 300 °C, a gradual mass shift to 6.8 MDa is observed. This progressive decrease in mass is attributed to combined loss of the viral envelope and partial loss of subunit(s). Notably, a concurrent increase in the abundance of feature at 82 kDa (assigned as prM-En) is observed (Figure 11B). This is accompanied by a pronounced broadening, affecting both this feature and the species at 172 kDa (assigned as [prM-En]_2_), with the FWHM increasing from ∼16 kDa to ∼40 kDa. While it is possible that some of these prM-En species arise from dissociation of [prM-En]_2_, the significant increase in FWHM may instead indicate that these species retain associated envelope components, such as lipids, and therefore originate from the 7.1 MDa VLPs.

**Figure 11.**
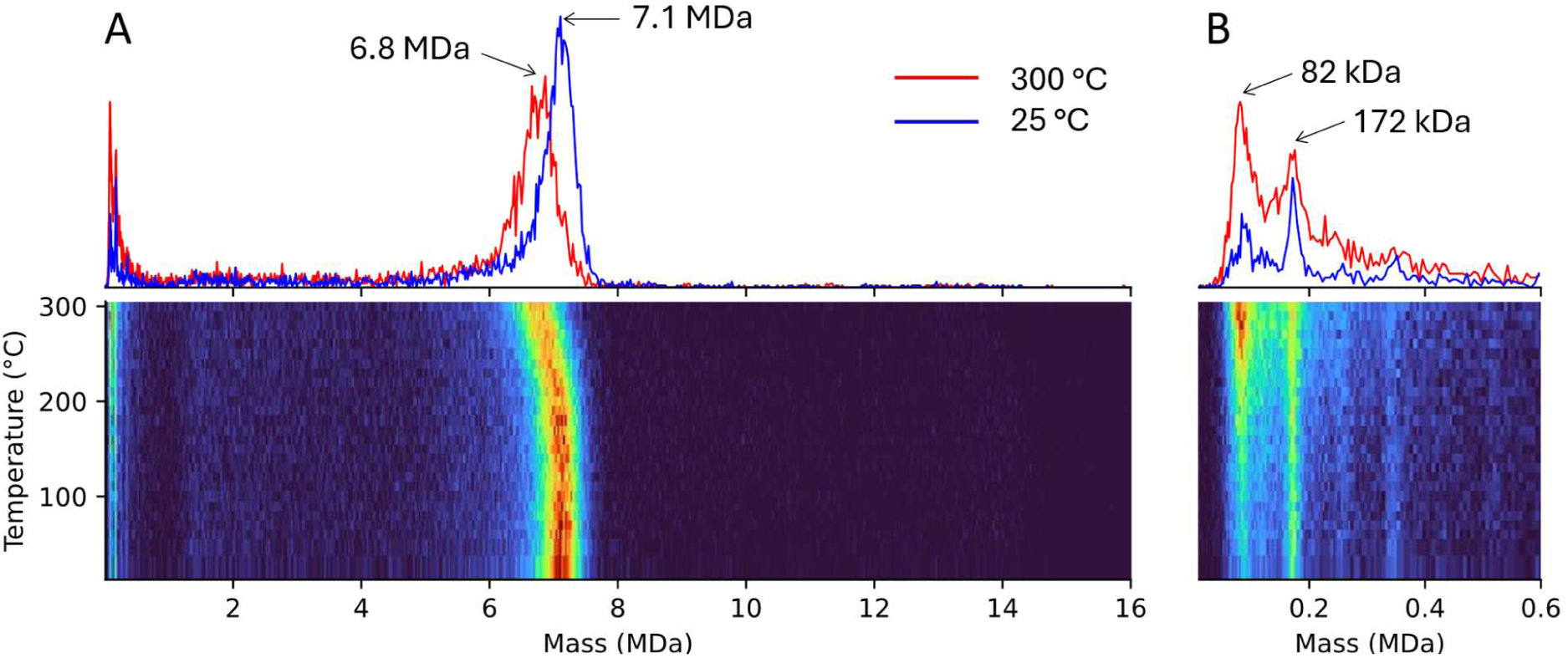
Activation of DENV VLPs using a heated-inlet interface. The panels show two-dimensional heatmaps of mass as a function of inlet tube temperature. A: entire mass range in which intact DENV VLP species are observed. B: low-mass region containing subunits and dissociation products. Data at each inlet temperature (in 10 °C increments) was acquired for 15 minutes.

Qualitatively similar behaviour was observed for other VLPs; for example, the CHIKV VLP data (“CHIKV VLPs fresh” and “CHIKV VLPs fresh inlet 300 degC” traces in interactive figure B, in SI) show a mass shift from 52.3 MDa to 46.6 MDa, accompanied by an apparent reduction in the charge state range, indicative of compaction of the envelope-less CHIKV VLP construct. The underlying patterns observed at the low-mass end for both CHIKV and DENV VLP species likely contain additional structural information (e.g., glycan or lipid heterogeneity) and warrant further investigation.

## Conclusions

The design and performance characteristics of a new benchtop CDMS instrument have been described. Beyond the ease-of-use, the key advantages of this new instrument stem from the combination of a wide *m/z* transmission range, detection efficiency and charge precision, enabling high-resolution mass measurements. Ion optics have been specifically optimized to cover a broad *m/z* range (up to *m/z* 250,000) with minimal tuning required. Detector response, governed by inductive sensing, is not mass dependent, making it particularly effective for observing large, highly charged particles, with hardware limit of approximately 3000 charges. The achievable *mass* resolution depends on charge precision and ranges from ∼20 to ∼160, owing to the instrument’s unique ability to reach the charge quantization threshold within practical trapping timescales (∼2 s). Although modest compared to the *m/z* resolution of traditional MS instruments, this corresponds to the highest *mass* resolution reported on a commercially available CDMS system to date.

In this work we have demonstrated the use of this novel instrument across several application areas, including protein complexes, AAV empty/full (E/F) ratio determination and large VLP characterization. While the characterization of vaccine and gene therapy products represents a primary focus, the instrument’s capabilities extend to other fields requiring accurate high-mass measurements. These include polymer science, glycobiology, nanoparticle characterization, and microplastics research, among others. Characterization in the 10-100 nm size range is challenging, with few good options between the molecular characterization tools used for molecules and the methods used to characterize bulk materials. The benchtop instrument described here helps to fill the void.

## Author Contributions

The manuscript was written through contributions of all authors.

## Notes

The authors declare the following competing financial interest(s): JU, CW, AS, MD, DE, DB, AH, KR, DL, AJ, and JB are employed by Waters Corporation who manufacture CDMS based instrumentation for sale. BD and MFJ are consultants to Waters Corporation. StepWave, XEVO and waters_connect are trademarks of Waters Corporation or its affiliates. PEEKsil is a trademark of Trajan Scientific and Medical. Orbitrap, Direct Mass Technology and Gibco are trademarks of Thermo Fisher Scientific Inc. Amicon and Milipore are trademarks of Merck KGaA. Bio-Rad and Micro Bio-Spin are trademarks of Bio-Rad Laboratories, Inc. Python is a trademark of Python Software Foundation. Hamilton is a trademark of Hamilton Medical Inc. PLOTLY is a trademark of Plotly Technologies Inc.

## ACKNOWLEDGMENT

The authors gratefully acknowledge the support and assistance of Prof. David Clemmer and the team at Megadalton Solutions (Daniel Botamanenko and Lohra Miller), which were fundamental to this work.

The authors also acknowledge the skill and expertise of the wider project team at Waters, whose contributions made this work possible: Ruchika Kogimtzis, Hong Ly, Adam Powell, Nicholas Smith, Duncan Leonard, Joseph Michienzi, Sidra Muntaha, Rosie Upton, Emilia Christofi, Rebecca D’Esposito, Lindsay Collins, Rob Lewis, Gareth Sunley, Jack McDonald, Alex Hughes, Danielle Foroughi, Chris Royle, Barry Dyson, Vicky Knox, David Ballantyne, Kate Yu, Martin Gill, and Stephen McDonald.

## REFERENCES

(1) Nollet, J. A. Essai Sur L’électricité Corps 1765.

(2) Dole, M.; Mack, L. L.; Hines, R. L.; Mobley, R. C.; Ferguson, L. D.; Alice, M. B. J. Chem. Phys. 1968, 49 (5), 2240–2249. 10.1063/1.1670391.

(3) Yamashita, M.; Fenn, J. B. J. Phys. Chem. 1984, 88 (20), 4451–4459. 10.1021/j150664a002.

(4) Fenn, J. B.; Mann, M.; Meng, C. K.; Wong, S. F.; Whitehouse, C. M. Science 1989, 246 (4926), 64–71. 10.1126/science.2675315.

(5) Alexandrov, M. L.; Gall, L. N.; Krasnov, N. V.; Nikolaev, V. I.; Pavlenko, V. A.; Shkurov, V. A. Dokl. Akad. Nauk SSSR 1984, 277, 379–383. 10.1002/rcm.3113.

(6) Chowdhury, S. K.; Katta, V.; Chait, B. T. J. Am. Chem. Soc. 1990, 112 (24), 9012–9013. 10.1021/ja00180a074.

(7) Ganem, B.; Li, Y. T.; Henion, J. D. J. Am. Chem. Soc. 1991, 113 (16), 6294–6296. 10.1021/ja00016a069.

(8) Katta, V.; Chait, B. T. J. Am. Chem. Soc. 1991, 113 (22), 8534–8535. 10.1021/ja00022a058.

(9) Baca, M.; Kent, S. B. H. J. Am. Chem. Soc. 1992, 114 (10), 3992–3993. 10.1021/ja00036a066.

(10) Ferrige, A. G.; Seddon, M. J.; Green, B. N.; Jarvis, S. A.; Skilling, J.; Staunton, J. Rapid Commun. Mass Spectrom. 1992, 6 (11), 707–711. 10.1002/rcm.1290061115.

(11) Skilling, J.; Richardson, K.; Brown, J.; Campuzano, I.; Green, B.; Wildgoose, J. In 58th ASMS Conference on Mass Spectrometry and Allied Topics; Salt Lake City, UT, 2010.

(12) Marty, M. T.; Baldwin, A. J.; Marklund, E. G.; Hochberg, G. K. A.; Benesch, J. L. P.; Robinson, C. V. Anal. Chem. 2015, 87 (8), 4370–4376. 10.1021/acs.analchem.5b00140.

(13) Wilm, M. S.; Mann, M. Int. J. Mass Spectrom. Ion Process. 1994, 136 (2), 167–180. 10.1016/0168-1176(94)04024-9.

(14) Wilm, M.; Mann, M. Anal. Chem. 1996, 68 (1), 1–8. 10.1021/ac9509519.

(15) Rostom, A. A.; Robinson, C. V. J. Am. Chem. Soc. 1999, 121 (19), 4718–4719. 10.1021/ja990238r.

(16) Harvey, S. R.; Gadkari, V. V.; Ruotolo, B. T.; Russell, D. H.; Wysocki, V. H.; Zhou, M. J. Am. Soc. Mass Spectrom. 2024, 35 (3), 646–652. 10.1021/jasms.3c00352.

(17) Tito, M. A.; Tars, K.; Valegard, K.; Hajdu, J.; Robinson, C. V. J. Am. Chem. Soc. 2000, 122 (14), 3550–3551. 10.1021/ja993740k.

(18) Uetrecht, C.; Versluis, C.; Watts, N. R.; Roos, W. H.; Wuite, G. J. L.; Wingfield, P. T.; Steven, A. C.; Heck, A. J. R. Proc. Natl. Acad. Sci. 2008, 105 (27), 9216–9220. 10.1073/pnas.0800406105.

(19) Ruotolo, B. T.; Benesch, J. L. P.; Sandercock, A. M.; Hyung, S.-J.; Robinson, C. V. Nat. Protoc. 2008, 3 (7), 1139–1152. 10.1038/nprot.2008.78.

(20) Heck, A. J. R. Nat. Methods 2008, 5 (11), 927–933. 10.1038/nmeth.1265.

(21) Chorev, D. S.; Baker, L. A.; Wu, D.; Beilsten-Edmands, V.; Rouse, S. L.; Zeev-Ben-Mordehai, T.; Jiko, C.; Samsudin, F.; Gerle, C.; Khalid, S.; Stewart, A. G.; Matthews, S. J.; Grünewald, K.; Robinson, C. V. Science 2018, 362 (6416), 829–834. 10.1126/science.aau0976.

(22) Konermann, L.; Ahadi, E.; Rodriguez, A. D.; Vahidi, S. Anal. Chem. 2013, 85 (1), 2–9. 10.1021/ac302789c.

(23) Wörner, T. P.; Bennett, A.; Habka, S.; Snijder, J.; Friese, O.; Powers, T.; Agbandje-McKenna, M.; Heck, A. J. R. Nat. Commun. 2021, 12 (1), 1642. 10.1038/s41467-021-21935-5.

(24) Yang, Y.; Ivanov, D. G.; Kaltashov, I. A. Anal. Bioanal. Chem. 2021, 413 (29), 7205–7214. 10.1007/s00216-021-03601-3.

(25) Köhler, G.; Milstein, C. Nature 1975, 256 (5517), 495–497. 10.1038/256495a0.

(26) Leavy, O. Nat. Immunol. 2016, 17 (1), S13–S13. 10.1038/ni.3608.

(27) Fu, Z.; Li, S.; Han, S.; Shi, C.; Zhang, Y. Signal Transduct. Target. Ther. 2022, 7 (1), 93. 10.1038/s41392-022-00947-7.

(28) Wesdemiotis, C.; Williams-Pavlantos, K. N.; Keating, A. R.; McGee, A. S.; Bochenek, C. Mass Spectrom. Rev. 2024, 43 (3), 427–476. 10.1002/mas.21844.

(29) Kohn, D. B.; Chen, Y. Y.; Spencer, M. J. Gene Ther. 2023, 30 (10), 738–746.10.1038/s41434-023-00390-5.

(30) Miesbach, W.; Meijer, K.; Coppens, M.; Kampmann, P.; Klamroth, R.; Schutgens, R.; Tangelder, M.; Castaman, G.; Schwäble, J.; Bonig, H.; Seifried, E.; Cattaneo, F.; Meyer, C.; Leebeek, F. W. G. Blood 2018, 131 (9), 1022–1031. 10.1182/blood-2017-09-804419.

(31) Day, J. W.; Mendell, J. R.; Burghes, A. H. M.; Olden, R. W. van; Adhikary, R. R.; Dilly, K. W. Mol. Ther. Methods Clin. Dev. 2023, 31. 10.1016/j.omtm.2023.101117.

(32) Burnham, B.; Nass, S.; Kong, E.; Mattingly, M.; Woodcock, D.; Song, A.; Wadsworth, S.; Cheng, S. H.; Scaria, A.; O’Riordan, C. R. Hum. Gene Ther. Methods Part B 2015, 26 (6), 228–242. 10.1089/hgtb.2015.048.

(33) Werle, A. K.; Powers, T. W.; Zobel, J. F.; Wappelhorst, C. N.; Jarrold, M. F.; Lyktey, N. A.; Sloan, C. D. K.; Wolf, A. J.; Adams-Hall, S.; Baldus, P.; Runnels, H. A. Mol. Ther. Methods Clin. Dev. 2021, 23, 254–262. 10.1016/j.omtm.2021.08.009.

(34) Young, G.; Hundt, N.; Cole, D.; Fineberg, A.; Andrecka, J.; Tyler, A.; Olerinyova, A.; Ansari, A.; Marklund, E. G.; Collier, M. P.; Chandler, S. A.; Tkachenko, O.; Allen, J.; Crispin, M.; Billington, N.; Takagi, Y.; Sellers, J. R.; Eichmann, C.; Selenko, P.; Frey, L.; Riek, R.; Galpin, M. R.; Struwe, W. B.; Benesch, J. L. P.; Kukura, P. Science 2018, 360 (6387), 423–427. 10.1126/science.aar5839.

(35) Asor, R.; Loewenthal, D.; Wee, R. van; Benesch, J. L. P.; Kukura, P. Annu. Rev. Biophys. 2025, 54 (Volume 54, 2025), 379–399. 10.1146/annurev-biophys-061824-111652.

(36) https://info.refeyn.com/l/983761/2025-02-10/53l85/983761/1739186792Bj5PnWlw/Refeyn_BioanalyticsWithMassPhotometry.pdf.

(37) Jarrold, M. F. Chem. Rev. 2022, 122 (8), 7415–7441. 10.1021/acs.chemrev.1c00377.

(38) Shockley, W. J. Appl. Phys. 1938, 9 (10), 635–636. 10.1063/1.1710367.

(39) Shelton, H.; Hendricks, C. D., Jr.; Wuerker, R. F. J. Appl. Phys. 1960, 31 (7), 1243–1246. 10.1063/1.1735813.

(40) Weinheimer, A. J. J. Atmospheric Ocean. Technol. 1988, 5 (2), 298–304. 10.1175/1520-0426(1988)005%253C0298:TCIOAC%253E2.0.CO;2.

(41) Fuerstenau, S. D.; Benner, W. H. Rapid Commun. Mass Spectrom. 1995, 9 (15), 1528–1538. 10.1002/rcm.1290091513.

(42) Fuerstenau, S. D.; Benner, W. H.; Thomas, J. J.; Brugidou, C.; Bothner, B.; Siuzdak, G. Angew. Chem. Int. Ed. 2001, 40 (3), 541–544. 10.1002/1521-3773(20010202)40:3%253C541::AID-ANIE541%253E3.0.CO;2-K.

(43) Benner, W. H. Anal. Chem. 1997, 69 (20), 4162–4168. 10.1021/ac970163e.

(44) Contino, N. C.; Jarrold, M. F. Int. J. Mass Spectrom. 2013, 345–347, 153–159. 10.1016/j.ijms.2012.07.010.

(45) Todd, A. R.; Alexander, A. W.; Jarrold, M. F. J. Am. Soc. Mass Spectrom. 2020, 31 (1), 146–154. 10.1021/jasms.9b00010.

(46) Keifer, D. Z.; Shinholt, D. L.; Jarrold, M. F. Anal. Chem. 2015, 87 (20), 10330–10337. 10.1021/acs.analchem.5b02324.

(47) Todd, A. R.; Jarrold, M. F. J. Am. Soc. Mass Spectrom. 2020, 31 (6), 1241–1248. 10.1021/jasms.0c00081.

(48) Parikh, R. A.; Alexander, A. W.; Jarrold, M. F. J. Am. Soc. Mass Spectrom. 2025, 36 (10), 2290–2298. 10.1021/jasms.5c00236.

(49) Gamero-Castaño, M. Rev. Sci. Instrum. 2007, 78 (4), 043301. 10.1063/1.2721408.

(50) Doussineau, T.; Kerleroux, M.; Dagany, X.; Clavier, C.; Barbaire, M.; Maurelli, J.; Antoine, R.; Dugourd, P. Rapid Commun. Mass Spectrom. 2011, 25 (5), 617–623. 10.1002/rcm.4900.

(51) Elliott, A. G.; Merenbloom, S. I.; Chakrabarty, S.; Williams, E. R. Int. J. Mass Spectrom. 2017, 414, 45–55. 10.1016/j.ijms.2017.01.007.

(52) Doussineau, T.; Yu Bao, C.; Clavier, C.; Dagany, X.; Kerleroux, M.; Antoine, R.; Dugourd, P. Rev. Sci. Instrum. 2011, 82 (8), 084104. 10.1063/1.3628667.

(53) Poschenrieder, W. P. Int. J. Mass Spectrom. Ion Phys. 1972, 9 (4), 357–373. 10.1016/0020-7381(72)80020-2.

(54) Barkhanskiy, A.; Cabrera, E.; Liggett, E.; Hoare, T.; Ishikawa, Y.; Chapman, R.; Hoyes, J.; Biggar, B.; Barran, P. In 73rd ASMS Conference on Mass Spectrometry and Allied Topics; Baltimore, MD, 2025.

(55) Draper, B. E.; Jarrold, M. F. J. Am. Soc. Mass Spectrom. 2019, 30 (6), 898–904. 10.1007/s13361-019-02172-z.

(56) Kafader, J. O.; Beu, S. C.; Early, B. P.; Melani, R. D.; Durbin, K. R.; Zabrouskov, V.; Makarov, A. A.; Maze, J. T.; Shinholt, D. L.; Yip, P. F.; Kelleher, N. L.; Compton, P. D.; Senko, M. W. J. Am. Soc. Mass Spectrom. 2019, 30 (11), 2200–2203. 10.1007/s13361-019-02309-0.

(57) Direct Mass Technology. https://www.thermofisher.com/uk/en/home/industrial/mass-spectrometry/liquid-chromatography-mass-spectrometry-lc-ms/lc-ms-systems/direct-mass-technology.html (accessed 2026-05-28).

(58) Smith, R. D.; Cheng, X.; Brace, J. E.; Hofstadler, S. A.; Anderson, G. A. Nature 1994, 369 (6476), 137–139. 10.1038/369137a0.

(59) Deslignière, E.; Yin, V. C.; Ebberink, E. H. T. M.; Rolland, A. D.; Barendregt, A.; Wörner, T. P.; Nagornov, K. O.; Kozhinov, A. N.; Fort, K. L.; Tsybin, Y. O.; Makarov, A. A.; Heck, A. J. R. Nat. Methods 2024, 21 (4), 619–622. 10.1038/s41592-024-02207-8.

(60) Ebberink, E. H. T. M.; Yin, V. C.; Deslignière, E.; Barendregt, A.; Wörner, T. P.; Fort, K. L.; Makarov, A. A.; Heck, A. J. R. J. Am. Chem. Soc. 2025, 147 (13), 10925–10934. 10.1021/jacs.4c13393.

(61) Bowen, K. P.; Liu, W.; Szabo, Z.; Senko, M. W. In 73rd ASMS Conference on Mass Spectrometry and Allied Topics; Baltimore, MD, 2025.

(62) Rayleigh, Lord. Lond. Edinb. Dublin Philos. Mag. J. Sci. 1882, 14 (87), 184–186. 10.1080/14786448208628425.

(63) Fernandez de la Mora, J. Anal. Chim. Acta 2000, 406 (1), 93–104. 10.1016/S0003-2670(99)00601-7.

(64) Keifer, D. Z.; Motwani, T.; Teschke, C. M.; Jarrold, M. F. J. Am. Soc. Mass Spectrom. 2016, 27 (6), 1028–1036. 10.1007/s13361-016-1362-8.

(65) Clemmer, D. E.; Jarrold, M. F. J. Mass Spectrom. 1997, 32 (6), 577–592. 10.1002/(SICI)1096-9888(199706)32:6%253C577::AID-JMS530%253E3.0.CO;2-4.

(66) Giles, K.; Gordon, D. 2010. https://www.waters.com/nextgen/us/en/library/application-notes/2010/A-New-Conjoined-RF-Ion-Guide-for-Enhanced-Ion-Transmission.html (accessed 2026-05-20).

(67) Green, M.; Wildgoose, J. L.; Pringle, S. D.; Giles, K. US7683314B2, March 23, 2010.

(68) Hogan, J. A.; Jarrold, M. F. J. Am. Soc. Mass Spectrom. 2018, 29 (10), 2086–2095. 10.1007/s13361-018-2007-x.

(69) Parikh, R. A.; Draper, B. E.; Jarrold, M. F. Anal. Chem. 2024, 96 (7), 3062–3069. 10.1021/acs.analchem.3c05087.

(70) Todd, A. R.; Jarrold, M. F. Anal. Chem. 2019, 91 (21), 14002–14008. 10.1021/acs.analchem.9b03586.

(71) Bertuccio, G.; Rehak, P.; Xi, D. Nucl. Instrum. Methods Phys. Res. Sect. Accel. Spectrometers Detect. Assoc. Equip. 1993, 326 (1), 71–76. 10.1016/0168-9002(93)90334-E.

(72) Todd, A. R.; Alexander, A. W.; Jarrold, M. F. J. Am. Soc. Mass Spectrom. 2020, 31 (1), 146–154. 10.1021/jasms.9b00010.

(73) Richardson, K.; Giles, K.; Haris, A.; Draper, B. E.; Jarrold, M. F. In 73rd ASMS Conference on Mass Spectrometry and Allied Topics; Baltimore, MD, 2025.

(74) Hunter, J. D. Comput. Sci. Eng. 2007, 9 (3), 90–95. 10.1109/MCSE.2007.55.

(75) 2015. https://plotly.com/python/.

(76) Ebberink, E. H. T. M.; Yin, V. C.; Deslignière, E.; Barendregt, A.; Wörner, T. P.; Fort, K. L.; Makarov, A. A.; Heck, A. J. R. J. Am. Chem. Soc. 2025, 147 (13), 10925–10934. 10.1021/jacs.4c13393.

(77) Kostyuchenko, V. A.; Zhang, Q.; Tan, J. L.; Ng, T.-S.; Lok, S.-M. J. Virol. 2013, 87 (13), 7700–7707. 10.1128/jvi.00197-13.

(78) Kuhn, R. J.; Zhang, W.; Rossmann, M. G.; Pletnev, S. V.; Corver, J.; Lenches, E.; Jones, C. T.; Mukhopadhyay, S.; Chipman, P. R.; Strauss, E. G.; Baker, T. S.; Strauss, J. H. Cell 2002, 108 (5), 717–725. 10.1016/S0092-8674(02)00660-8.

(79) Johnson, A.; Dodes Traian, M.; Walsh, R. M.; Jenni, S.; Harrison, S. C. J. Virol. 99 (2), e01809–24. 10.1128/jvi.01809-24.

(80) Metz, S. W.; Thomas, A.; White, L.; Stoops, M.; Corten, M.; Hannemann, H.; de Silva, A. M. Virol. J. 2018, 15 (1), 60. 10.1186/s12985-018-0970-2.

