## Supplementary material for "A Charge Detection Mass Spectrometer for the Analysis of Megadalton-sized Molecules": CDMS Supporting Information.pdf

### Additional details on High Duty Cyle (HDC) mode

While travelling from the accumulation region to the ELIT, ions separate according to their mass-to-charge dependent times-of-flight (ToFs):

$$ToF = k\sqrt{m/z}$$

Therefore, opening of the ELIT trap needs to be delayed such that it will trap only the ions of interest. The range of *m/z* values that can be captured using this approach can be estimated by adding the ELIT residence time (i.e., 1.75T) of the ion, to its ToF:

$$ToF_{heavier} = k\sqrt{m/z} + 1.75\sqrt{\frac{m/z}{C}}$$

Where constants k and C are voltage and geometry dependent. For the instrument presented here, the ratio of *m/z* values corresponding to lighter and heavier ions that can be simultaneously trapped, will take a form of:

$$\frac{\left(k\sqrt{C} + 1.75\right)^2}{k^2C} \approx 1.16$$

While this method can achieve near 100% duty cycle for a small *m/z* range, it is not suitable for applications requiring uniform enhancement over a broad *m/z* range, such as e.g. full-empty AAV ratio measurement (*m/z* range 20k-40k).

### Calibration of CDMS analyser

#### Calibration of charge

Careful amplifier design and appropriate signal processing ensures that the magnitude *A* of the fundamental peak corresponding to an ion in the FT is nearly proportional to its charge state. In practice a linear calibration is employed to allow for (and measure) small offsets in charge measurement:

$$z = c_1 + c_2A$$

These constants are determined using protein standards that can be resolved by *m/z* in order that a representative magnitude can be determined for each charge state. Because the charge states are resolved by *m/z*, a charge calibration can be formed by determining representative magnitudes corresponding to individual charge state peaks for a number of standards. Having done this, the parameters $c_1$ and $c_2$ can be determined using a straightforward linear fit.

#### Calibration of *m/z*

It is desirable to utilize calibration standards having a wide range of charge states to establish a high-quality charge calibration. The same standards can also contribute usefully to *m/z* calibrations. However higher mass standards can also carry a significant amount of extra mass in the form of adducts. This uncertain additional mass must be taken into account during *m/z* calibration.

As has already been stated in the main text, the relationship between frequency and *m/z* in an electrostatic ion trap is given by:

$$\frac{m}{z} = \frac{C}{f^2}$$

Where *C* is a constant (to be calibrated) which depends on the geometry of the trap, ion energy, and the applied voltages. The calibration dataset will typically consist of series of data for *N* standards having known masses $m_1, m_2, \ldots m_N$. Each of these standards will be represented in the data by a number of charge states $z=z_{n1}, z_{n2},\ldots$ for the *i*'th standard ($1 \leq i \leq N$). We allow that each standard has some *a-priori* unknown amount of excess mass $\delta_i$ attached which we currently assume to be the same for each charge state so that the observed *m/z* values $d_{ij}$ (where j indexes charge states), obtained using a provisional calibration will be:

$$d_{ij} = g\frac{m_i + \delta_{ij}}{z_{ij}} \pm \sigma_{ij}$$

where *g* is an *a-priori* unknown scaling factor ($g=1.0$ if the provisional calibration constant *C* is exactly correct). Following calibration, *C* will be updated to a revised value $C'=C/g$. The $\sigma_{ij}$ *are* uncertainties in the *m/z* measurements which are obtained by peak detecting a spectrum such as that shown in Figure S 8B. Note that for simplicity we have ignored the mass of charge carriers here: although relatively small these are easily included in the analysis.

We adopt a Bayesian approach, which requires that we assign prior probability distributions to the unknown parameters *g* and $\delta_i$. To avoid unnecessary complication we assign a uniform prior to the calibration parameter *g*. Because the excess mass parameters $\delta_i$ are known to be positive, an exponential prior for these is appropriate:

$$\Pr(\delta_i) = \exp -\frac{\delta_i}{\Lambda_i}$$

Where $\Lambda_i$ is the expected scale of additional mass (e.g. 1% of $m_i$). Assuming a Gaussian likelihood function, the joint probability of the data and parameters is:

$$\Pr(d, \delta_1, \delta_2, \ldots, g) = \prod_i \frac{1}{\Lambda_i} \exp\left(-\frac{\delta_i}{\Lambda_i}\right) \prod_{j \in z_i} \frac{1}{\sqrt{2\pi}\sigma_{ij}} \exp\left(-\frac{\left(d_{ij} - g\frac{m_i+\delta_i}{z_{ij}}\right)^2}{2\sigma_{ij}^2}\right)$$

$$= \frac{1}{(2\pi)^{N/2}} \prod_{ij} \frac{1}{\sigma_{ij}} \prod_i \frac{1}{\Lambda_i} \exp\left(-\frac{(\delta_i - \delta_{i0})^2}{2S_i^2}\right) \exp\left(\sum_i \left(\frac{B_i^2}{4A_i} - C_i\right)\right)$$

Where in the second line we have simply rearranged the expression (to facilitate further analysis) and defined the new symbols:

$$A_i = \frac{1}{2} g^2 X_i \qquad B_i = g^2 m_i X_i - g Y_i + \frac{1}{\Lambda_i}$$

$$C_i = \frac{1}{2} g^2 m_i^2 X_i - g m_i Y_i + \frac{1}{2} Z_i \qquad S_i = \frac{1}{\sqrt{2A_i}} \qquad \delta_{i0} = -\frac{B_i}{2A_i}$$

$$X_i = \sum_{j \in z_i} \frac{1}{z_j^2 \sigma_{ij}^2} \qquad Y_i = \sum_{j \in z_i} \frac{d_{ij}}{z_j \sigma_{ij}^2} \qquad Z_i = \sum_{j \in z_i} \frac{d_{ij}^2}{\sigma_{ij}^2}$$

Since we are not primarily interested in the additional mass parameters $\delta_i$, they can be "marginalized" i.e. integrated out of the full joint probability to give the joint probability of $g$ and the data:

$$\Pr(d, g) = \int_0^\infty \int_0^\infty \Pr(d, \delta_1, \delta_2, \ldots, g)\, \mathrm{d}\delta_1 \mathrm{d}\delta_2 \ldots$$
$$\propto \exp\left(\sum_i \left(\frac{B_i^2}{4A_i} - C_i\right)\right) \prod_i S_i \left(1 + \operatorname{erf}\left(\frac{\delta_{i0}}{\sqrt{2}S_i}\right)\right)$$

Where erf is the error function. Everything in this joint probability distribution is known except $g$, so despite the apparent complexity it is just a 1D distribution. It is therefore straightforward to obtain samples of $g$ to determine the required calibration scaling $g$ together with the associated uncertainty.

### Description of and use of the interactive figures

In order to enable readers to interrogate the data presented here, we have included two interactive figures. These are provided as two .html files, which should be opened in an internet browser (“Interactive Figure A – mz vs charge.html” and “Interactive Figure B – mass vs charge.html”). Each contains a mass or *m/z* histogram located at the top and the corresponding scatter plot at the bottom. By default, all data series are displayed; however, they can be selected or deselected by clicking on the respective entries in the legend. Double-left-clicking on a legend entry removes all other series. The interactive plots support zoom and pan functionality. Zooming is enabled by left-clicking and dragging over the area of interest in either the scatter or histogram plots. To autoscale, double-left-click on either plot. Pan functionality can be enabled by clicking on the button in top right corner.

Interactive figures include the following datasets:

| ***Legend entry*** | ***Description*** |
|---|---|
| CHIKV VLPs incubated | Chikungunya VLP, 0.3 mg/ml in 200 mM ammonium acetate solution + 0.01 % P-188, incubated for a week at room temperature prior to analysis. 100 ms trapping time. |
| CHIKV VLPs fresh inlet 300 degC | As above, analysed immediately after buffer exchange with inlet temperature of 300 °C. 100 ms trapping time. |
| CHIKV VLPs fresh | As above, analysed immediately after buffer exchange with inlet temperature of 25 °C. 100 ms trapping time. |
| DENV VLPs | Dengue serotype 1 VLP, 0.3 mg/ml in 200 mM ammonium acetate solution. |
| DENV VLPs inlet 300 degC | Dengue serotype 1 VLP, 0.3 mg/ml in 200 mM ammonium acetate solution, inlet temperature of 300 °C. 100 ms trapping time. |
| AAV8 E-F mixture | AAV8 empty and full mixture (50/50); data from figure 8 (10 injections combined). |
| Beta-Galactosidase 100ms | β-Galactosidase, 0.2 mg/ml in 200 mM ammonium acetate solution, 100 ms trapping time. |
| Beta-Galactosidase 2000ms | As above, 2000 ms trapping time. |
| Glutamate Dehydrogenase | Glutamate dehydrogenase, 0.2 mg/ml in 200 mM ammonium acetate solution. 100 ms trapping time. |
| Monoclonal Antibody | Waters Intact mAb Mass Check Standard, 0.01 mg/ml in 200 mM ammonium acetate solution. 100 ms trapping time. |
| Enolase | Enolase, 0.2 mg/ml in 200 mM ammonium acetate solution. 100 ms trapping time. |
| Myoglobin denatured | Myoglobin, 0.025 mg/ml in water and 0.1% formic acid. 100 ms trapping time. |

Note that, to construct histograms of all datasets on a common bin grid, logarithmically spaced bins were used. Data presented in the main manuscript were instead binned using fixed-width bins; consequently, peak heights may appear slightly different from those shown in the main figures.

### Additional figures

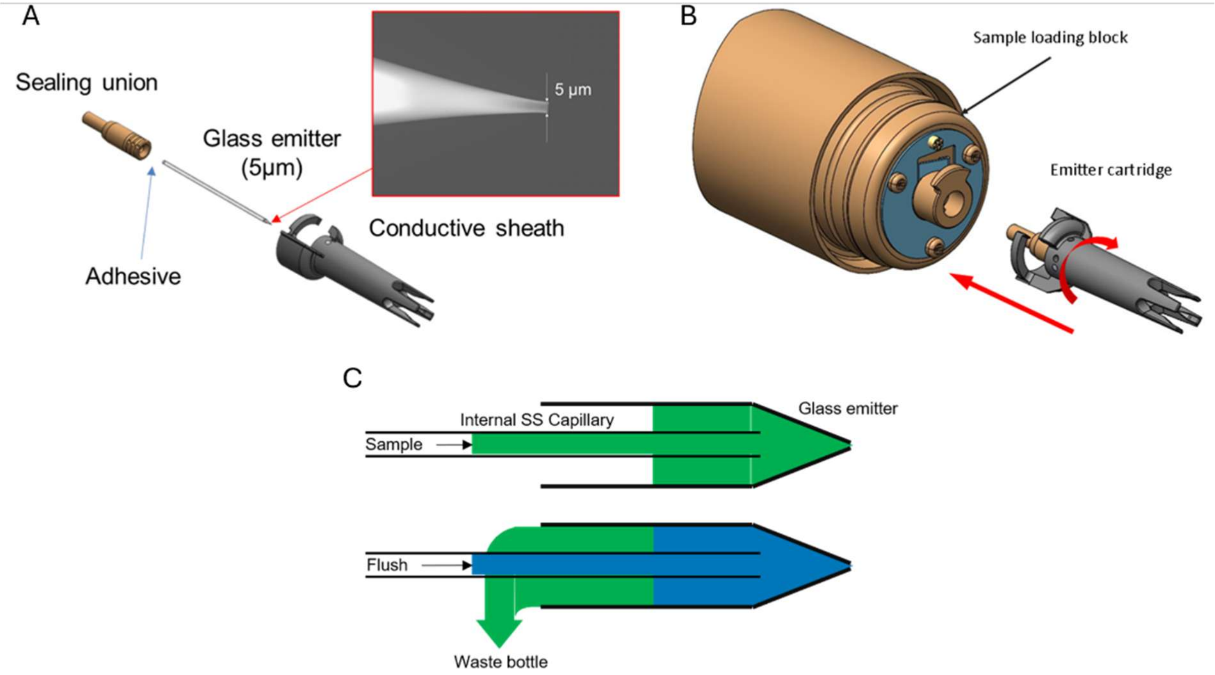

Figure S 1. Detailed view of ion source components: (A) Emitter cartridge design; (B) Emitter cartridge replacement; (C) Schematic of the glass emitter and internal SS capillary enabling sample delivery and flush operation.

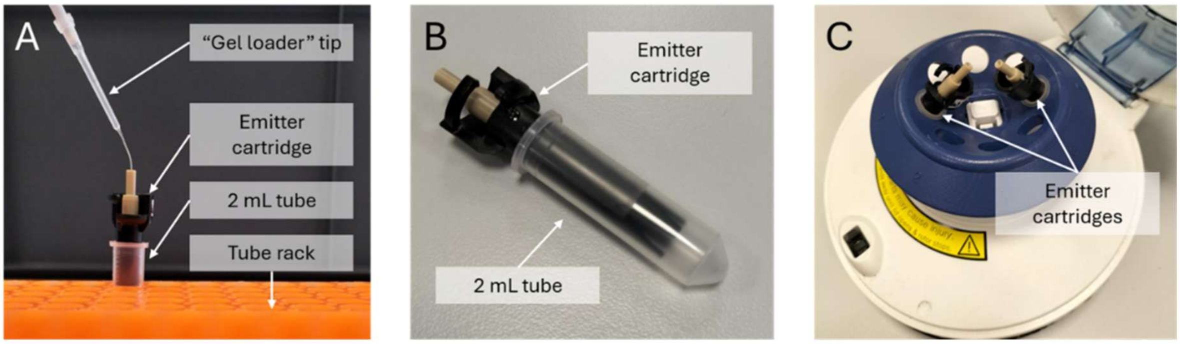

Figure S 2. Offline workflow for low-volume samples: (A) Direct loading of the sample into the emitter cartridge using a standard pipette with a gel-loading tip; (B) Storage of the emitter cartridge in a standard 2 mL test tube; (C) Brief "spin-down" in a mini-centrifuge to ensure the sample reaches the tip of the glass emitter.

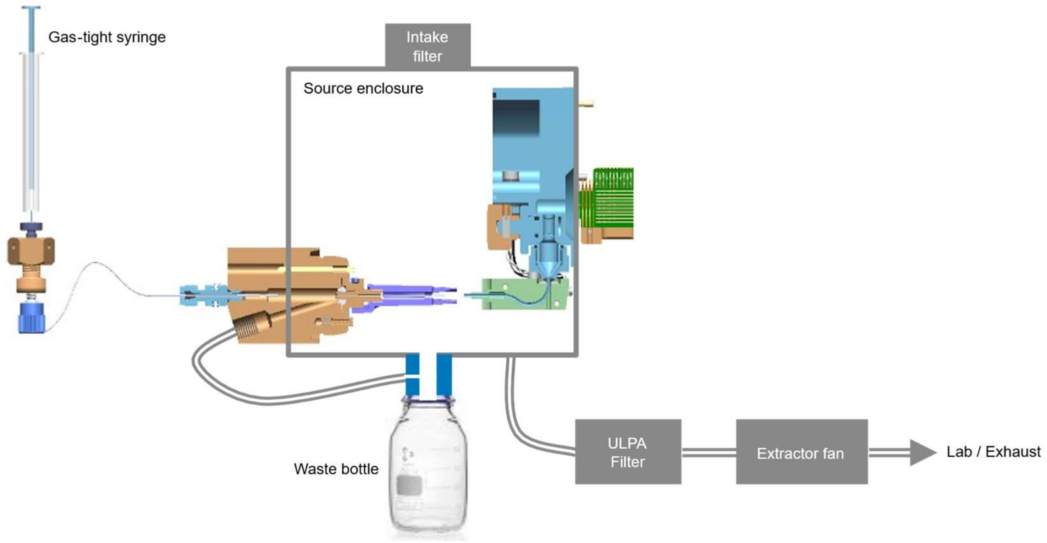

Figure S 3. Schematic of nESI source enclosure and extraction system.

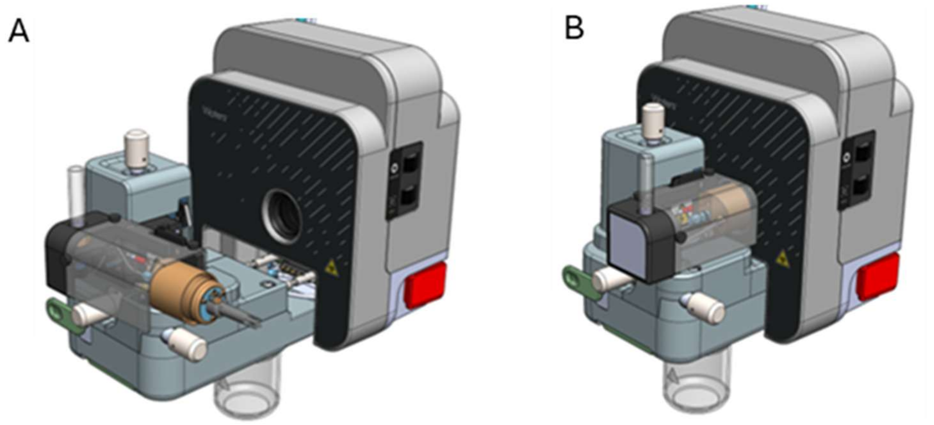

Figure S 4. External design of the nESI source enables easy access and emitter cartridge replacement: (A) The retracted sprayer manifold can be rotated by 90° to facilitate emitter cartridge replacement; (B) Operational configuration with the manifold reinserted.

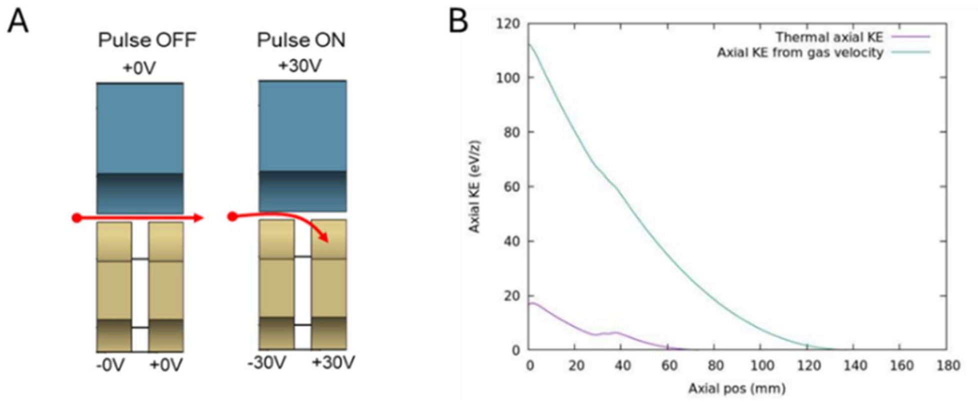

Figure S 5. Beam attenuation and cooling: (A) Beam attenuator switching states. (B) Simulation of the axial kinetic energy (KE) of a 50 MDa particle carrying 400 charges. The horizontal axis corresponds to the axial position along the length of Segmented Quadrupole 1 (Seg Quad 1), where 0 mm corresponds to Aperture 1, ~38 mm to the position of the beam attenuator electrodes and 180 to Aperture 2. The initial axial KE (at position 0 mm) was approximated using two models: thermal motion of gas molecules and gas velocity derived from pressure-driven flow.

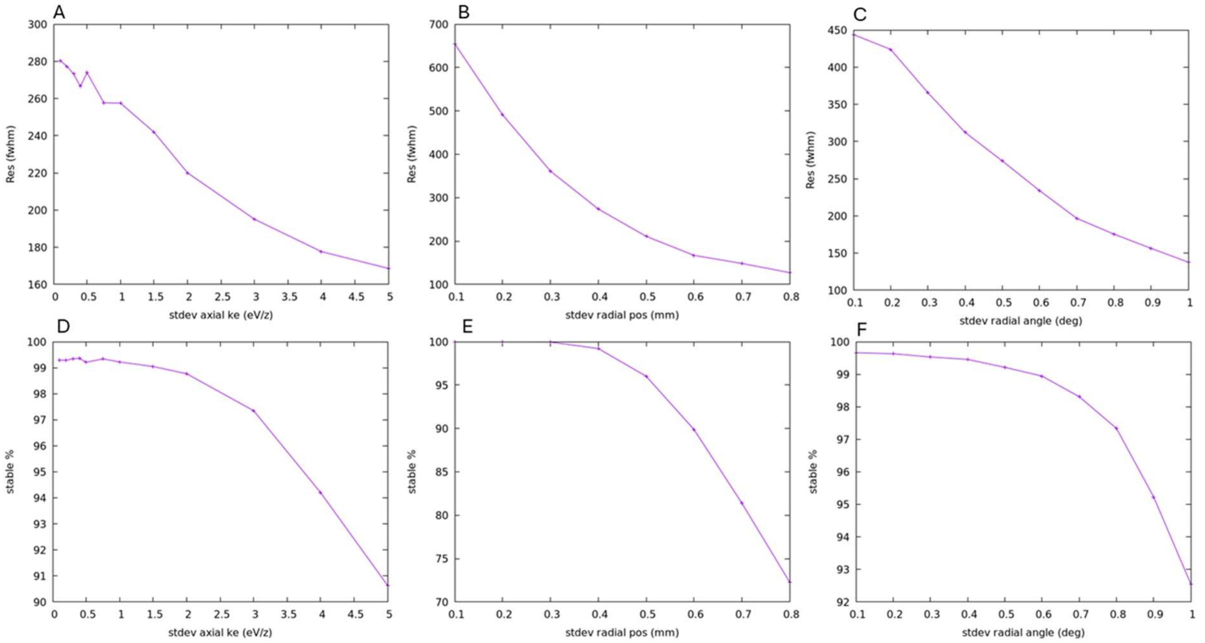

Figure S 6. A: Simulations of *m/z* resolution and stability in dimensionally perfect, 2D ELIT model vs ion beam phase space. Top row panels demonstrate *m/z* resolution vs standard deviation in axial kinetic energy (A), radial position (B) and beam divergence angle (C). Bottom row panels demonstrate number of ions that survive the first 200 passes (no significant further losses are observed in the ideal model device) vs standard deviation in axial kinetic energy (D), radial position (E) and beam divergence angle (F).

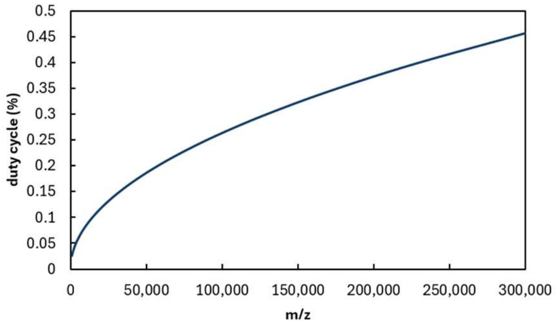

Figure S 7. Calculated ELIT CDMS duty cycle vs *m/z* in Random Trapping Mode.

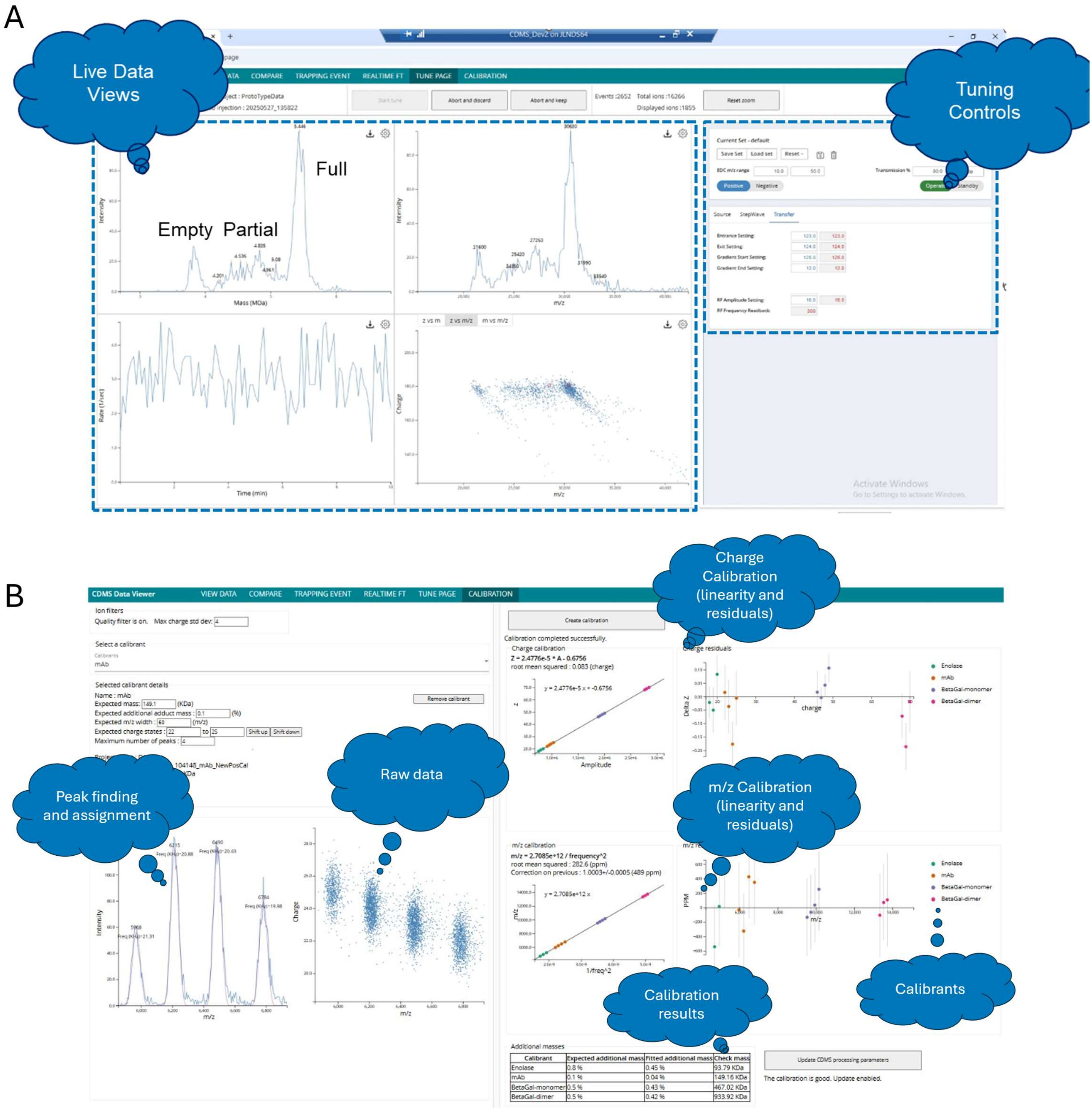

Figure S 8. Instrument control software: (A) Live data views and tuning controls; (B) Calibration page and example calibration dataset. Calibrant spectra (Enolase, Monoclonal Antibody and β-Galactosidase) are included in interactive figures A and B.

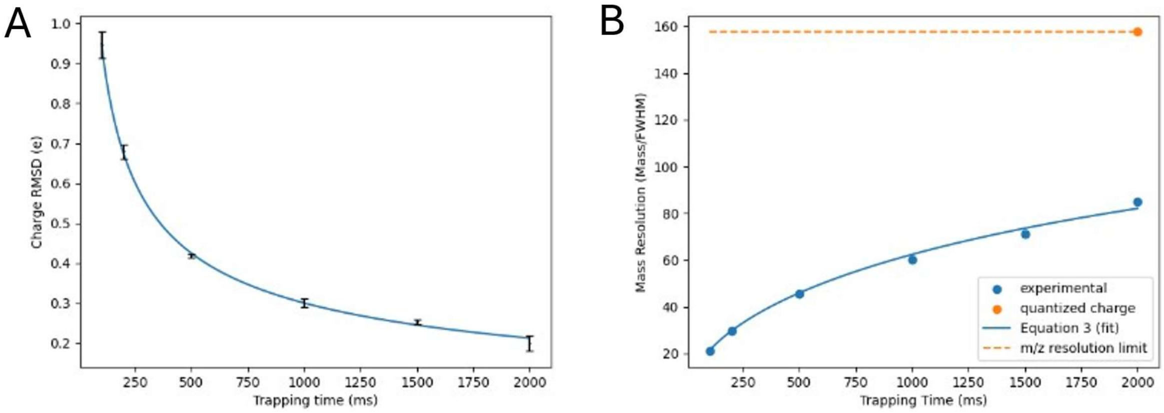

Figure S 9. Charge and mass precision as a function of trapping time measured for β-galactosidase ions: (A) Charge RMSD vs. trapping time; points represent experimental data, and the fitted line corresponds to Equation (4) in the main manuscript; (B) Mass resolution as a function of trapping time; points represent experimental data, and the fitted line corresponds to Equation (3) in the main manuscript, with $\sigma_z$ term set to $\sigma_{z\_ref}$ /sqrt(trapping_time/$t_{ref}$), where $\sigma_{z\_ref} = 0.93$ $e$ and $t_{ref} = 100$ ms.

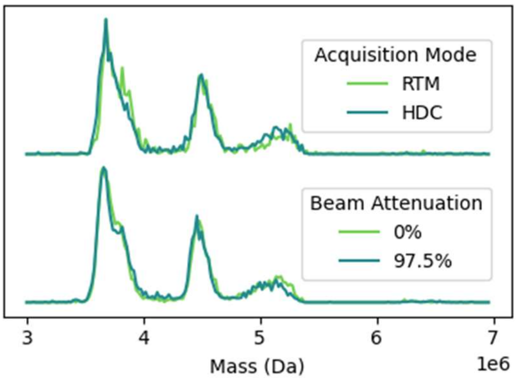

Figure S 10. AAV mass distributions are not strongly influenced by concentration, acquisition mode and beam attenuation: (A) Mass spectra of empty and full AAV8 mixtures prepared at concentration of $1\times10^{10}$ vp/mL and analysed using RTM and HDC mode; (B) the corresponding mixtures prepared at concentration of $1\times10^{12}$ vp/mL and using different beam attenuation settings.
